# Wildfires drive trade-offs in ammonia-oxidizing groups and promote denitrification to increase soil emissions of nitric oxide (NO) and nitrous oxide (N_2_O) in California chaparral

**DOI:** 10.64898/2026.09.10.749748

**Authors:** Elizah Z. Stephens, Alexander H. Krichels, Aral C. Greene, Sharon Zhao, Chloe Reid, Jamie Irby, Maria E. Ordoñez, Dominika Lewicka-Szczebak, Erin J. Hanan, Sydney I. Glassman, Peter M. Homyak

**Author notes:** **Corresponding author:** Elizah Z. Stephens1, Department of Environmental Sciences, University of California, Riverside CA 92521, USA.

## Abstract

Wildfires can promote trade-offs in soil nitrifier and denitrifier communities that affect post-fire nitrogen (N) cycling and emissions of nitric oxide (NO), nitrous oxide (N_2_O), and dinitrogen (N_2_). For example, by increasing soil pH and ammonium (NH_4_^+^), wildfires could increase the abundance of ammonia-oxidizing bacteria (AOB) over archaea (AOA). Because AOB and AOA process N differently, nitrifier trade-offs may affect N emissions and nitrate (NO_3_^-^) supply with downstream effects on denitrifier activity, leading us to ask: Do trade-offs between AOA and AOB abundance and shifts in denitrifying processes influence N emissions over time after wildfire? We selectively inhibited AOA and AOB communities from soil collected over four seasonal time points one year before and after a chaparral wildfire and used stable isotopes to parse denitrifier contributions to N_2_O emissions. Over one year after the wildfire, soil pH increased from 6.1 to 7.0, soil extractable NH_4_^+^ increased 30-fold, NO_3_^-^ increased 5-fold, and NO_2_^-^ increased 9-fold. AOB *amoA* gene copy numbers increased 20-fold one year after fire, while AOA abundance remained unchanged. Post-fire soil NO emissions increased 63-fold over one year, with varied contributions from all nitrifier groups. Soil N_2_O emissions peaked eight months after fire (2271 ± 634 ng N_2_O-N g^-1^ soil), with increased contributions from AOA. Wildfire increased δ^15^N^SP^_N2O_ and δ^15^N^bulk^_N2O_ values, suggesting increased N_2_O reduction to N_2_. Overall, wildfire increased AOB abundance relative to AOA, promoting nitrification activity and providing intermediates to denitrifiers to increase emissions of NO and N_2_O for up to one year after fire.

## 1. Introduction

As global changes in climate and land use interact to increase wildfire extent and/or severity (Cunningham et al., 2024; Ellis et al., 2022; Williams et al., 2019), ecosystem cycling of essential nutrients like nitrogen (N) may become increasingly altered. Wildfires can drive immediate net ecosystem N loss during combustion of organic matter and N losses can continue as fires modify soils and microbial communities that produce N gases such as nitric oxide (NO), nitrous oxide (N_2_O), and dinitrogen (N_2_; Stephens & Homyak, 2023). For example, increased soil pH and N-rich ash left behind after fire (Neary et al., 1999) may favor ammonia-oxidizing bacteria (AOB) over ammonia-oxidizing archaea (AOA; Ball et al., 2010; Long et al., 2014). Since AOB are often associated with higher NO and N_2_O emissions than AOA (Prosser et al., 2019), the increase in AOB could accelerate nitrification rates and increase emissions. Enhanced nitrification activity may also provide nitrate (NO_3_^-^) to denitrifiers spanning a broad range of taxa, which may undergo turnover after fire (Pulido-Chavez et al., 2022), further altering NO and N_2_O emissions patterns. Not only could changes in NO and N_2_O emissions contribute to ecosystem N loss, but NO is an air pollutant at high concentrations and a precursor for tropospheric ozone formation (Crutzen, 1979; Ostro et al., 2006) and N_2_O is a powerful greenhouse gas and a major contributor to stratospheric ozone depletion (Griffis et al., 2017). Thus, understanding the functional trade-offs that underly N loss and emissions after wildfire contributes to our ability to predict ecosystem recovery trajectories, greenhouse gas emissions, and regional air quality as wildfire activity increases globally.

NO and N_2_O emissions in soils are primarily produced by microbes capable of nitrification and denitrification. During nitrification, ammonia (NH_3_; measured in soils as ammonium; NH_4_^+^) is oxidized to nitrite (NO_2_^-^) and NO_3_^-^ aerobically, releasing NO and N_2_O as by-products (Firestone & Davidson, 1989). During denitrification, NO_3_^-^ is sequentially reduced to NO_2_^-^, NO, N_2_O, and N_2_ under low oxygen availability (Knowles, 1982). Major controls on nitrification and denitrification rates in undisturbed soils have traditionally emphasized factors such as N substrate availability (NH_4_^+^ for nitrification and NO_3_^-^ for denitrification), pH, and soil moisture (Firestone and Davidson, 1989), which control how N “leaks” from soil microbial pathways as gaseous byproducts. These factors have proven useful predictors of soil N gas emissions in most undisturbed contexts (Davidson et al., 2000), but wildfires also cause major microbial community and functional turnover (Krichels et al., 2025; Pulido-Chavez et al., 2022; Pulido Barriga et al., 2025; Stephens et al., 2026), making post-fire contexts more difficult to predict (Stephens et al., 2026; Stephens & Homyak, 2023). Similar to Grime’s C-S-R (Competitor-Stress-Ruderal) model developed for plant community succession after disturbance (Grime, 1977), post-fire microbial communities may exhibit trade-offs for post-fire resource acquisition, thermotolerance, and fast growth (Enright et al., 2022), with important implications for N cycling (Pulido Barriga et al., 2025). Therefore, considering post-fire trade-offs in the functions of nitrifying and denitrifying microbial groups over time alongside abiotic changes in N substrates, pH, and water content may improve prediction of post-fire N gas emission variability.

Nitrification is rate-limited by the activity of organisms which oxidize NH_3_ to NO_2_^-^ and NO_3_^-^ such as chemoautotrophic ammonia-oxidizing archaea in phylum Thaumarchaeota (AOA) and ammonia-oxidizing bacteria in genus *Nitrosomonas* and *Nitrosococcus* (AOB; Carey et al., 2016; Prosser et al., 2019). Some heterotrophic microbes are also capable of ammonia-oxidation; however, the major groups of organisms thought to contribute to bulk nitrification in soils are autotrophic (Barraclough & Puri, 1995; Robertson & Groffman, 2006). While performing similar functions, bacterial and archaeal ammonia-oxidizers operate within different soil niches and emit N gases at different rates (Prosser et al., 2019). For example, AOA thrive in acidic environments with low NH_3_ availability and are associated with lower NO and N_2_O emissions relative to higher-emitting AOB, which tend to favor environments with higher pH and more abundant NH_3_ availability (Carey et al., 2016; Hatzenpichler, 2012; Prosser et al., 2019). Because wildfire can increase soil NH_4_^+^ availability and pH (Hanan et al., 2017; Smithwick et al., 2005; Ulery et al., 2017), post-fire soil conditions may promote a trade-off between nitrifier groups, favoring AOB dominance. AOB have been associated with up to 50% higher emissions of N_2_O relative to AOA (Prosser et al., 2019) and may contribute the bulk of NO emissions from some forest soils (Mushinski et al., 2019). Therefore, by increasing soil pH and NH_4_^+^ to favor AOB abundance over AOA (Avrahami & Bohannan, 2009; Ball et al., 2010; Long et al., 2014), wildfire may enhance post-fire soil NO and N_2_O emissions (**Figure 1**).

**Figure 1.**
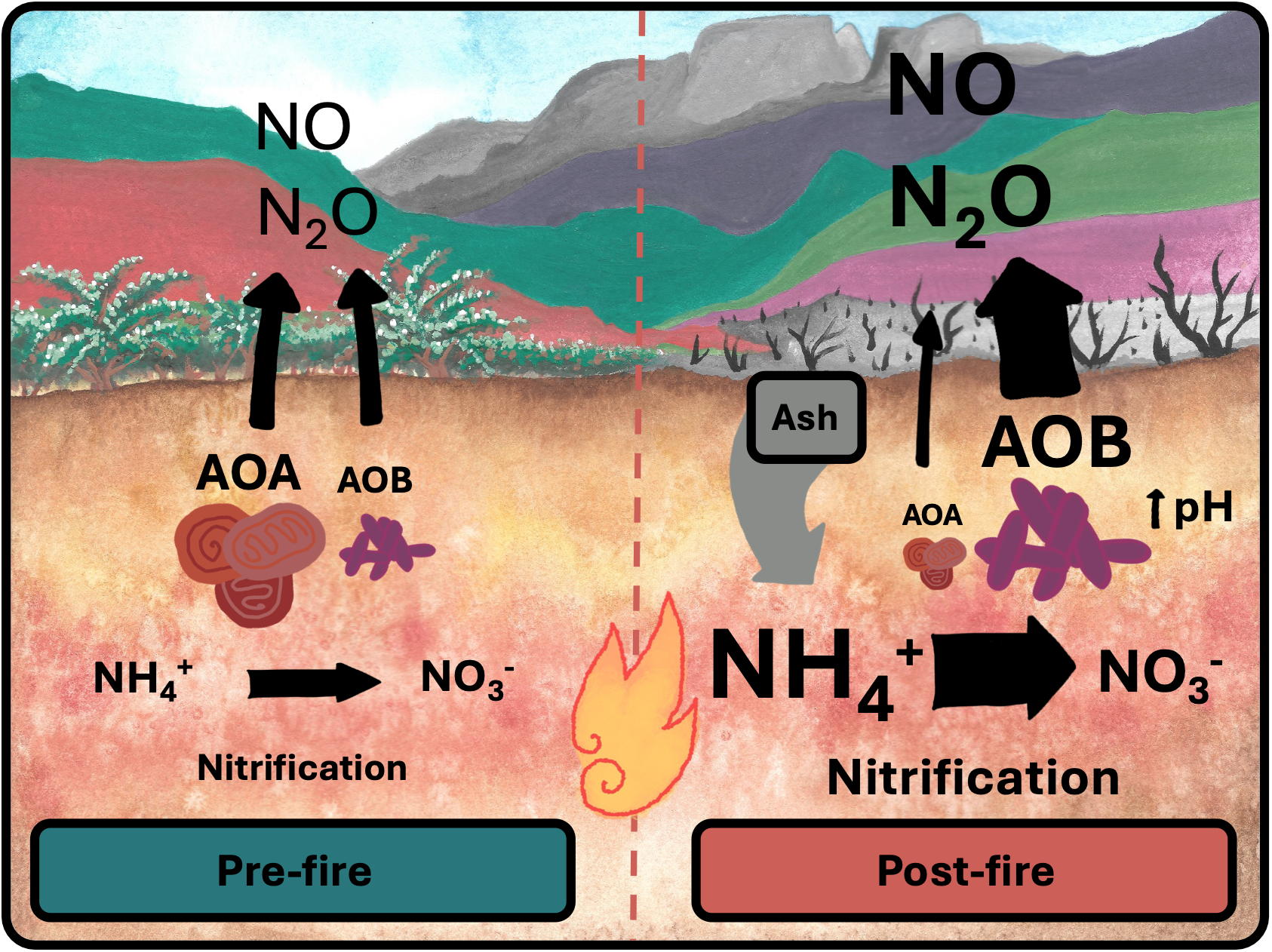
Hypothesized pre- and post-fire N-cycling changes where sizes of arrows and boxes represent expected changes in N pools, process rates, trace N gas fluxes, and communities of ammonia-oxidizing archaea (AOA) and ammonia-oxidizing bacteria (AOB). We expect wildfire ash to increase soil NH_4_^+^ availability and pH, stimulating nitrification rates and increasing AOB activity to support larger NO and N_2_O emissions.

While nitrification is carried out by a relatively narrow range of taxa, denitrification can be performed by heterotrophic bacteria (Hayatsu et al., 2008); by some nitrifiers such as AOB which can carry out nitrifier denitrification (Stein, 2019; Wrage et al., 2001); and partially by fungi (N_2_O is the final product as fungi are not known to perform further reduction to N_2_; Aldossari & Ishii, 2021). Increased NO_3_^-^ from nitrification after fire may stimulate denitrification activity when soils become water-saturated and suboxic during heavy rains, favoring increased NO, N_2_O, and N_2_ emissions following fire (Dannenmann et al., 2018; Stephens & Homyak, 2023). However, the stimulatory effect of wildfire on denitrification may vary over time as groups of denitrifying organisms with different resource acquisition strategies trade off after wildfire. For example, fungal communities may be reduced to a greater extent and have slower recovery times than bacteria (Glassman et al., 2023; Pressler et al., 2019; Pulido-Chavez et al., 2022); therefore we might expect to see changes in the contributions of denitrifier groups to NO and N_2_O emissions. Predicted increases in AOB abundance may also increase nitrifier denitrification, as AOB are capable of nitrifier denitrification while AOA are not (Stein, 2019). Relatively recent advances in isotope techniques make it possible to distinguish the major groups involved in N_2_O production (bacteria, nitrifier denitrification, fungi) by using isotopocules of N_2_O (δ^15^N^bulk^_N2O_, δ^18^O_N2O_, and site preference δ^15^N^SP^_N2O_, which reflects the placement of ^15^N in the central α and peripheral β positions in the N_2_O molecule; Yu et al., 2020). Because distinct enzymatic pathways used by different denitrifying organisms produce unique isotopic fractionation patterns, these isotopic techniques can provide insight into how fires might drive shifts in the major microbial groups involved in N_2_O production. We therefore ask: how might post-fire trade-offs between AOA and AOB nitrification and shifts in denitrifying processes shape soil N cycling over time to influence gaseous N losses?

A unique opportunity to address this question arose when the KNP Complex fire burned a chaparral watershed in Sequoia National Park, CA, USA, in September 2021, where we had conducted seasonal (fall, winter, spring, summer) measurements of soil nutrients and contributions of AOA and AOB to NO and N_2_O emissions before the fire. We continued this sampling scheme for one year after the fire to directly compare the biogeochemical functions pre- and post-wildfire. Dry shrublands, like this chaparral site, are increasingly fire-prone (Baeza et al., 2005; Ellis et al., 2022; Park et al., 2021; Syphard et al., 2023), and have been shown to respond strongly to wildfire by increasing NO and N_2_O emissions (Stephens et al., 2026; Stephens & Homyak, 2023), making this site an ideal location to address our question. We hypothesized that fire would increase AOB abundance relative to AOA due to high soil pH and NH_4_^+^ availability (**H1a**), thereby increasing AOB-derived NO and N_2_O emissions (**H1b; Figure 1)**. We also expected elevated post-fire nitrification activity to increase NO_3_^-^ availability to denitrifiers and promote soil N_2_O emissions with isotopic signatures associated with changing contributions of microbial groups that trade off after fire (**H2**). We tested these hypotheses by 1) quantifying AOA and AOB *amoA* gene abundances with quantitative PCR, 2) comparing the contributions of AOA and AOB to NO and N_2_O fluxes using selective inhibitors, and 3) characterizing the isotopic composition of N_2_O to detect changes in soil denitrification processes at four seasonal timepoints over one year pre- and post-fire. Overall, we assess how post-fire trade-offs of nitrifying and denitrifying communities contribute to the loss of N as NO and N_2_O from a chaparral ecosystem.

## 2. Methods

### 2.1 Site description

In September 2021, the Chamise Creek watershed in Sequoia National Park (36° 30’ 47” N, 118 ° 48’ 26” W) burned in the KNP Complex wildfire. As the site has been an active long-term biogeochemical monitoring site since 2014 with pre-fire records of N pools (Homyak et al., 2014) and AOA/AOB activity (Krichels et al., 2024), this unplanned wildfire created a unique opportunity to compare N cycling in pre- and post-fire soils. Prior to the fire, the site was a mature chamise (*Adenostoma fasciculatum)* dominated chaparral shrubland in a 4.2 ha watershed that had not burned since 1960 (Li et al., 2006). The site is at 700 m elevation with 10 – 17 % slope that drains into a single catchment (Chamise Creek). Climate at the site is Mediterranean with cool, wet winters and dry, hot summers, and receives ∼670 mm of precipitation from November-April with average monthly air temperatures ranging from 8 to 40 °C. Pre-fire, soil bulk density was ∼1.2 g cm^-3^ in the upper 10 cm (Homyak et al., 2014) and soils were classified as coarse-loamy, mixed, superactive, thermic Ultic Haploxeralfs in the Ashmountain series with 17% clay, 63% sand, and 20% silt (**Table S1**). Burn severity of vegetation at the site was classified as moderate by the US Geological Survey Burned Area Reflectance Classification (https://burnseverity.cr.usgs.gov/viewer/?product=BAER; **Figure S1**).

### 2.2 Soil sampling and monitoring

Soils were collected before and after fire with a 7.5 cm diameter auger from the top 10 cm of mineral soil (A horizon; litter and ash were removed if present and auger was cleaned with ethanol between plots) across a 50 m transect with plots at 10 m intervals (n = 5; **Figure S1**). Pre-fire samples were collected in fall (October 19, 2020), winter (February 9, 2021), spring (May 6, 2021), and summer (August 27, 2021). One month after the fire was extinguished, we resumed this sampling scheme to collect post-fire soils in fall (October 24, 2021), winter (February 7, 2022), spring (May 19, 2022), and summer (September 1, 2022; **Table S2**).

Soil cores were transported to the University of California Riverside and sieved (4 mm) within 72 h post-sampling. Subsamples were immediately set aside to be stored at −20 °C for DNA extraction and qPCR analysis (section 2.5), and to oven-dry to measure soil water content (∼10 g). Soils were stored at 4 °C for no more than 48 h before they were mixed with 2 M KCl to measure soil extractable NH_4_^+^, NO_2_^-^, and NO_3_^-^ concentrations (Hanan et al., 2016). Soil extractable NO_2_^-^ was measured with nanopure water because KCl underestimates nitrite concentrations (Homyak et al., 2015). Soil extracts were analyzed colorimetrically at the Environmental Sciences Research Laboratory (https://envisci.ucr.edu/research/environmental-sciences-research-laboratory-esrl) at UCR for NH_4_^+^ [SEAL method Environmental Protection Agency (EPA)-126-A], NO_3_^-^ (SEAL method EPA-129-A), and NO_2_^-^ (SEAL method EPA-137-A) using a SEAL AQ-2 discrete analyzer. Soil gas flux incubations with nitrifier inhibitions (section 2.3) were performed on soils stored at 4 °C for no longer than 1 week following sample collection. Remaining soils were stored for isotopic analyses by air-drying at 25°C. Soil pH was measured in a 1:2 solution of air-dried soil in deionized water with a pH meter. Bulk soil total C and N and δ^13^C and ^15^N were measured with an elemental analyzer (Costech Elemental Combustion System) coupled with an isotope ratio mass spectrometer (Delta V Advantage IRMS, δ^15^N precision = 0.06 ‰) at the UC Riverside Facility for Isotope Ratio Mass Spectrometry (FIRMS; https://ccb.ucr.edu/facilities/firms). Water holding capacity (WHC) was calculated as the amount of water a saturated soil could hold after freely draining against gravity in a sealed container to minimize water losses to evaporation for 48 h.

### 2.3 qPCR of amoA genes

Subsamples of homogenized soil cores (n = 5 at each seasonal sampling which had been collected over an entire year pre-fire as well as post-fire, or n = 40 total) were stored at −20°C within 72 h of soil sampling until soil DNA was extracted from 0.25 g of soil with Qiagen DNeasy _PowerSoil_ Pro Kits as previously published (Joukhajian et al., 2026; Krichels et al., 2024). After DNA extraction, quantitative polymerase chain reaction (qPCR) was used to quantify the number of bacterial and archeal *amoA* genes as a proxy for AOA and AOB abundance (Beman et al., 2008). Bacterial *amoA* genes were targeted using the AmoA1F/amoA2R primer set (Rotthauwe et al., 1997) and the Arch-amoAF/ArchamoAR primer set was used for archaeal *amoA* (Francis et al., 2005). Triplicate reactions containing 5 μL of 2X master mix (Forget-Me-Not _EvaGreen_ qPCR Master Mix, Biotium, Inc., Fremont, CA), 0.8 μL of 25 mM MgCl2, 0.25 μL of 2 mg mL^-1^ bovine serum albumin, 0.125 μL of 20 μM forward and reverse primer, 2.5 μL ddH_2_O, and 1.2 μL sample DNA were run for each qPCR (Krichels et al., 2024) on the Bio-Rad CFX Opus 384 Real-Time PCR System (Bio-Rad, Hercules, CA). The following protocol was used to amplify bacterial *amo*A: 5 min at 95 °C, followed by 40 cycles of 45 s at 95 °C, 30 s at 56 °C and 60 s at 72 °C, and for archaeal *amoA*: 4 min at 95 °C, followed by 40 cycles of 30 s at 95 °C, 45 s at 53 °C and 60 s at 72 °C (Krichels et al 2024). We used serial dilutions of the amoA gene of *Nitrosomonas europaea* ATCC, 19718 as a standard for bacterial *amoA* and Crenarchaeota genomic fragment 54d9 as a standard for archaeal *amoA* (Beman et al., 2008; Eberwein et al., 2020; Francis et al., 2005). We calculated gene copy numbers per g soil as copy number = (starting concentration (ng/ μL) * (1 μL in reaction) * 6.022 * 10^^23^ (Avogadro’s constant)) / (length of standard (bp) * 660 ng (average weight of a base pair)* 10^^9^ (average number of cells per g of soil; Joukhajian et al., 2026)).

### 2.4 Selective inhibitions of AOA and AOB

To understand trade-offs between AOA- and AOB-driven nitrification, we used soil microcosms treated with selective inhibitors 1-octyne (to inactivate AOB ammonia oxidation; Taylor et al., 2013) and acetylene (to inactivate both AOB and AOA ammonia oxidation) to find the difference in NO and N_2_O emissions from each group following methods in Mushinski et al., (2019). We were able to conduct only one seasonal sampling before the unplanned wildfire burned at our study site, so n = 5 at each timepoint for pre-fire fall and post-fire winter, spring, summer, and fall for n = 25 total. Briefly, soil cores were homogenized and three 50 g soil subsamples were sealed in airtight jars (120 mL) equipped with rubber septa. The first replicate was injected with 4 μmol L^-1^ 1-octyne to inhibit AOB ammonia-oxidation, the second was injected with 6 μmol L^-1^ acetylene to inhibit total autotrophic nitrification (both AOA and AOB ammonia-oxidation), and the third was an unaltered control. These treatments were incubated for 24 h at 25 °C prior to measurement of NO and N_2_O. To inhibit bacterial or archaeal growth during the inhibition incubations, we made solutions of growth inhibitors mixed with DI water containing either 220 μg g^-1^ soil kanamyacin for the AOB inhibition treatment, or 220 μg g^-1^ soil kanamyacin + 800 μg g^-1^ soil fusidic acid + 200 μg g^-1^ soil nitrapyrin for the total nitrifier treatment. At the beginning of the measurement, antibiotic growth inhibitor solutions or control solutions of just DI water were added to wet the soils and bring them up to 100% WHC, which was chosen to approximate a heavy rainfall event which can saturate soils and stimulate microbial and abiotic processes involved in both NO and N_2_O production (Birch, 1958; Eberwein et al., 2020). NO and N_2_O fluxes from soils treated with inhibitors were continuously measured for 72 h as described in section 2.4. Fluxes associated with each inhibition treatment were used to calculate the individual contributions of AOA and AOB nitrification to NO and N_2_O fluxes (Krichels et al., 2024; Mushinski et al., 2019). The activity of AOB was estimated by subtracting the 1-octyne AOB inhibition treatment from the control; the activity of AOA was calculated by subtracting the acetylene total nitrifier inhibition treatment from the 1-octyne AOB inhibition, and the “Other” category (includes processes such as heterotrophic nitrification, abiotic, and denitrification) was considered equal to the acetylene total nitrifier inhibition treatment. Given these low concentrations of acetylene, we do not expect interferences with N_2_O production from denitrification and N_2_O reduction to N_2_ (Smith et al., 1978).

Soil extractions of NH_4_ ^+^ or NO_3_^-^ were performed on soils pre- and post-72 h gas flux incubations to calculate net N transformation rates as the change in NH_4_ ^+^ or NO_3_^-^ per day simultaneously with soil gas fluxes. Net N transformation rates were calculated as the difference in combined NH_4_^+^ and NO_3_^-^ (net N mineralization rate) or NO_3_^-^ (net nitrification rate) between final and initial measurements over 72 h.

### 2.5 Soil fluxes of NO and N_2_O

Soil NO and N_2_O fluxes were measured for each inhibition treatment (n = 5 at each timepoint: fall pre-fire and winter, spring, summer, and fall post-fire; n = 25 total) by multiplexing the jars to a continuous flow trace gas analyzer system as described in detail by Krichels et al. (2024). This consists of a recirculating sample loop controlled by a multiplexer (LI-8150, LI-COR Biosciences) connected to a laser N_2_O analyzer (Los Gatos Research, Inc.; Model 914–0027), an infrared CO_2_/H_2_O analyzer (LI-8100, LI-COR Biosciences), and a chemiluminescent NO_2_ analyzer (Scintrex LMA-3D, Unisearch Associates, Canada) fitted with a CrO_3_ converter to oxidize NO to NO_2_. When the CrO_3_ converter was removed, NO_2_ was not detectable, thus we assume our measurements were mostly NO. Each jar was measured continuously for 19 minutes at intervals of 2 h. Because NO is consumed by the NO_2_ analyzer, we used a 3-way solenoid valve to automatically route air from the recirculating sample loop at a rate of 1.5 mL min^-1^ into the NO_2_ analyzer halfway through the total 19-minute measurement cycle. At the same time, 1.5 mL min^-1^ zero-grade air (Ultra Grade Zero Air, Airgas, Radnor) was allowed to flow into the jars to replace the sample and prevent a vacuum. NO concentrations were allowed to equilibrate with the sample loop for 10 minutes and the last 30 s were averaged. Soil NO emissions were calculated as:

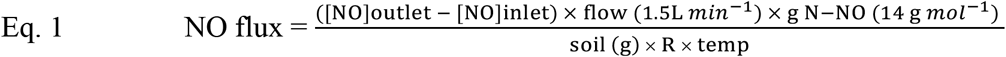

where [NO]outlet is the concentration of NO leaving the jar headspace (ppb), [NO]inlet is the concentration of NO entering the jar (assumed to be 0 ppb), soil (g) is the mass of soil in the jar (50 g), R is the molar gas constant (0.0821 L atm K^-1^ mol^-1^), and temp is the room air temperature (∼25°C). At the end of the 19-minute measurement cycle, the sample loop was purged for 1 minute with ambient air before the next jar was measured.

N_2_O emissions were calculated as the increase in concentration over the first 9 minutes of the measurement cycle when the NO_2_ analyzer was not connected using a publicly available R script (Andrews & Krichels, 2022). To prevent capturing gas fluxes as inhibitors might have been wearing off, the last 12 h of each 72-h incubation was trimmed and cumulative N_2_O and NO emissions were calculated using trapezoidal integration (trapZ function in R; R Core Team, 2024) over the first 48 h of the incubation period after wetting. Not every inhibitor replicate produced a decrease in NO or N_2_O emissions, resulting in negative fluxes for some groups. We report these and note that inhibitors produced the majority of negative values in burned soils, indicating a possible interference with biochar-like compounds such as woodchar or pyrogenic carbon (Pokharel & Chang, 2021). Water content was tracked by weighing every 24 h.

### 2.6 Natural abundance isotopic composition of N_2_O and NO

To better understand underlying processes contributing to N_2_O emissions, we analyzed the isotopic composition of N_2_O: δ^15^N^bulk^_N2O_ and δ^18^O_N2O_, where δ = ((Rsample/Rstandard)-1)×1000) in units of permil (‰) and R = ^15^N/^14^N or ^18^O/^16^O). We also measured site preference (δ^15^N^SP^_N2O_; which reflects the placement of ^15^N in the central (α) and peripheral (β) positions in the N_2_O molecule) as δ^15^N^SP^_N2O_ = δ^15^N^α^ − δ^15^N^β^. Subsamples (50 g) of air-dried soils were selected to represent summer, spring, fall, and winter for the year before the fire (n = 3 per season; 12 total) to reflect pre-fire processes. All post-fire soils sampled at each season were used (n = 5 for each season; 20 total) in anticipation of high variation of N_2_O production which has been observed in post-fire soils (Stephens & Homyak, 2023). Since there was little drying in the sealed jars compared to our continuous flow set-up (methods section 2.5), soils were wet up to 75 % WHC, which corresponded to water contents at peak N_2_O production during AOA/AOB incubations. Wetted soils were immediately sealed in air-tight 250 mL mason jars fitted with 1 L gas-tight bags (Cali-5-Bond; Calibrated Instruments LCC) pre-filled with a mixture of zero-grade air and 1 ppm N_2_O (Airgas part no. X02AI99C33A01B7) to ensure a high enough N_2_O concentration to allow for isotope characterization. The jar headspace was flushed with the same 1 ppm N_2_O mixture for 10 minutes then sealed and incubated at 25 °C for 24 h. In addition to allowing 24 h for the jar headspace to diffuse into the 1 L bag via an open 4-way stopcock (Calibrated Instruments, LCC), we ensured even mixing by pumping with a 50 mL syringe 10 times before sealing and removing the bag for analysis. N_2_O was analyzed within one week on a cavity ringdown infrared N_2_O analyzer (Los Gatos Research, Inc.) fitted with a Nafion water trap (PD-200 T-12MPS, Perma Pure LLC), a CO_2_ trap (Carbosorb, Elemental Microanalysis), and an activated charcoal and silica gel trap to remove excess water and volatile organic compounds (Sigma-Aldrich), which can interfere with N_2_O isotopomer measurements (scrubber set-up described by Stuchiner et al., 2021; as published in Krichels et al., 2023). Gas samples were passed through the scrubbers and withdrawn from the 1 L bag into the isotopic N_2_O analyzer at a rate of 80 mL min^-1^ to allow ∼10 minutes of measurement time where N_2_O concentrations and isotopomers were recorded every second. Because it took 5-6 minutes for N_2_O concentration to stabilize, we averaged the last ∼3 minutes to obtain N_2_O concentrations and δ^15^N^bulk^_N2O_, δ^18^O_N2O_, and δ^15^N^SP^_N2O_ values. These data were referenced to international reference materials from the United States Geological Survey USGS 51, 52, 32, 34, and 35 (USGS Reston Stable Isotope Laboratory) by applying individual standard curves (R^2^ > 0.99; concentration range 1-7.5 ppm N_2_O) for each isotopocule of N_2_O (^14^N^15^N^16^O, ^15^N^14^N^16^O, and ^14^N^14^N^18^O), isotopocule concentrations were converted to delta notation using the following equations:

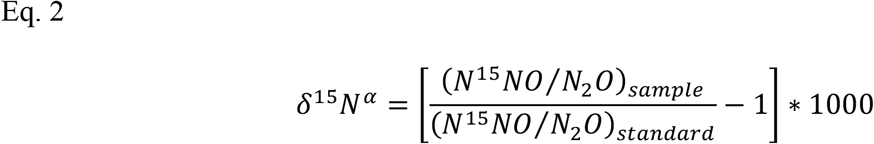

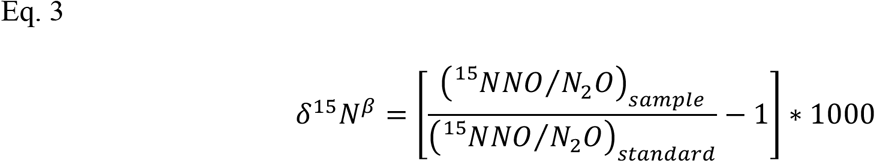

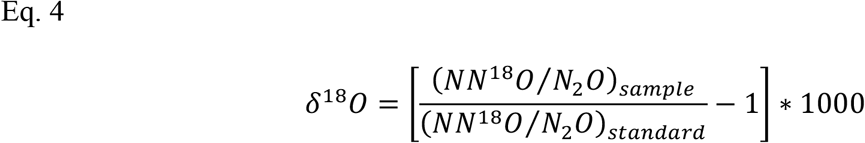

as reported by Krichels et al. (2023) and derived from Stuchiner et al., (2021). Averages over 3 minutes of 1s recoded values (n = 180) yielded ∼± <0.0001 standard deviations for all isotopocules concentrations in the range of our samples (1.6–2.8 ppm N_2_O; **Table S3**). To obtain the isotopic composition of soil-emitted N_2_O from the 1ppm N_2_O headspace we used a two-endmember mixing model:

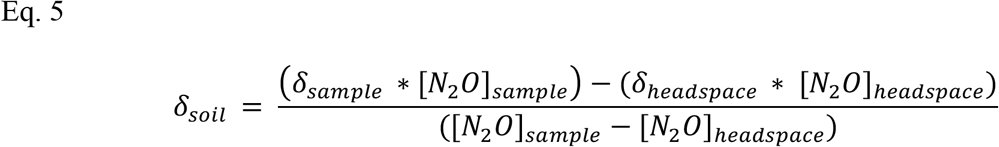

Where “[N_2_O]” represents the concentration of N_2_O in ppm, “soil” is the portion derived from soil only, “sample” is the total soil + headspace, and “headspace” is the isotopic composition and N_2_O concentration from the tank of 1 ppm N_2_O balanced with zero-grade air.

To compare our samples to literature-derived ranges for nitrification (Ni), nitrifier denitrification (nD), bacterial denitrification (bD), and fungal denitrification (fD; Lewicki et al., 2022; Yu et al., 2020), we adjusted for the δ^15^N-NO_3_^-^ and δ^15^N-NO_2_^-^ (substrates assumed to contribute to bacterial and fungal denitrification processes), δ^15^N-NH_4_^+^ (substrate assumed to contribute to nitrification and nitrifier denitrification) measured via acid trap diffusion (Irby et al., 2026) and adjusted for δ^18^O_H2O_ of the DI water added to incubations according to recommendations by Yu et al. (2020; **Methods S1; Table S4**). We then used the publicly available Isotope Fractionation And Mixing Evaluation (FRAME) model to partition fractional contributions of N_2_O production processes (Ni, fD, nD, bD) and the residual N_2_O fraction (r) to estimate the proportion of N_2_O reduction (**Table S5**; Lewicka-szczebak et al., 2020; Lewicki et al., 2022; Well & Flessa, 2010; Yu et al., 2020). We calculated cumulative N_2_ emissions by dividing the concentration of N_2_O at the end of the 48 h incubation by the estimated residual N_2_O fraction (Buchen et al., 2018; Lewicka-Szczebak, 2018). Fractionation processes contributing to N_2_O reduction were based on experimental data from Well & Flessa (2010) and model structure and auxiliary inputs matched example 5.4 in Lewicki et al. (2022).

To characterize the isotopic composition of NO emitted from soils, jar lids were fitted with NO_x_ precoated collection pads (Ogawa & Co., Pompano Beach, FL, USA) to characterize δ^15^N-NO and δ^18^O-NO as described in Homyak et al. (2016), see **Methods S2**.

### 2.7 Statistical analyses

All analyses were performed in R (R core team). To account for repeated soil sampling over time after fire, linear mixed effects models (LME; lme4 package in R; Bates et al. 2015) with a random effect for soil core replicate and an autocorrelation term to account for repeated measures over time that specified an autoregressive correlation structure (Zuur et al., 2009, https://m-clark.github.io/mixed-models-with-R/) were used to test for differences before and after fire. To determine differences between pre- and post-fire measurements for each season (**H1a, H2**), variables were modeled with an interaction between burn status and season as the predictor variables and fitted with Restricted Maximum Likelihood Estimation (REML). If there was an overall significant effect, then post-hoc analysis was performed using the “emmeans” package in R (Lenth 2023) to determine differences between burned and unburned in each season. To compare post-fire AOA and AOB-derived gas fluxes and N transformation rates to before the fire (**H1b**; only one season was available pre-fire), the same model was used without the interaction term for season to reference each post-fire timepoint to the pre-fire measurements. If the fire × treatment interaction was significant, a pairwise comparison of each treatment to pre-fire baseline was applied with treatment-wise contrasts (trt.vs.ctrl method with the Holm correction in emmeans). We used a Shapiro-Wilks test to verify that each model met normality assumptions (Shapiro & Wilk, 1965), and when assumptions were not met, a log-transformation was applied (as was the case for cumulative NO, N_2_O, and gene copy numbers for AOA and AOB).

## 3. Results

### 3.1 Wildfire altered soil properties and N cycling

Wildfire increased soil extractable NH_4_^+^ compared to pre-fire measurements in every season of our study (**Figure 2A**), representing a 30-fold average increase over one-year post-fire (average total post-fire increase of 26.1 ∝g NH_4_^+^-N g soil^-1^ over the pre-fire year; **Table S1**; fire: p <0.001). Soil extractable NO_3_^-^ increased 5-fold (post-fire increase of 6.9 ∝g NO_3_^-^-N g soil^-1^) across the entire year post-fire (**Table S1**; fire × season p <0.001), driven by a peak in summer (summer increase of 28.2 ∝g NO_3_^-^-N g soil^-1^ above pre-fire; **Figure 2B**; post-hoc summer p <0.001). The 9-fold post-fire increase in soil extractable NO_2_^-^ (average post-fire increase of 0.6 ∝g NO_2_^-^-N g soil^-1^; **Table S1**; fire × season: p < 0.001) was also driven by a single peak of 1.8 ∝g NO_2_^-^-N g soil^-1^ in post-fire spring (post-hoc spring p < 0.001; **Figure 2C**). Soil pH increased on average from 6.1 to 7.0 after fire, remaining significantly elevated compared to before the fire for all seasons (**Figure 2D**; **Table S1**; fire p = 0.009). Wildfire increased bulk soil C by 31% (average post-fire increase of 9 mg C g soil^-1^; fire: p <0.001), and bulk soil N by 29% (increase of 0.5 mg N g soil^-1^; fire: p <0.001), over one year after fire. Bulk soil δ^15^N was not affected by fire (fire p = 0.3), however, δ^13^C increased after fire by 0.9 ‰ and remained enriched across all seasons compared to before the fire (fire × season: p < 0.001; **Table S1**).

**Figure 2.**
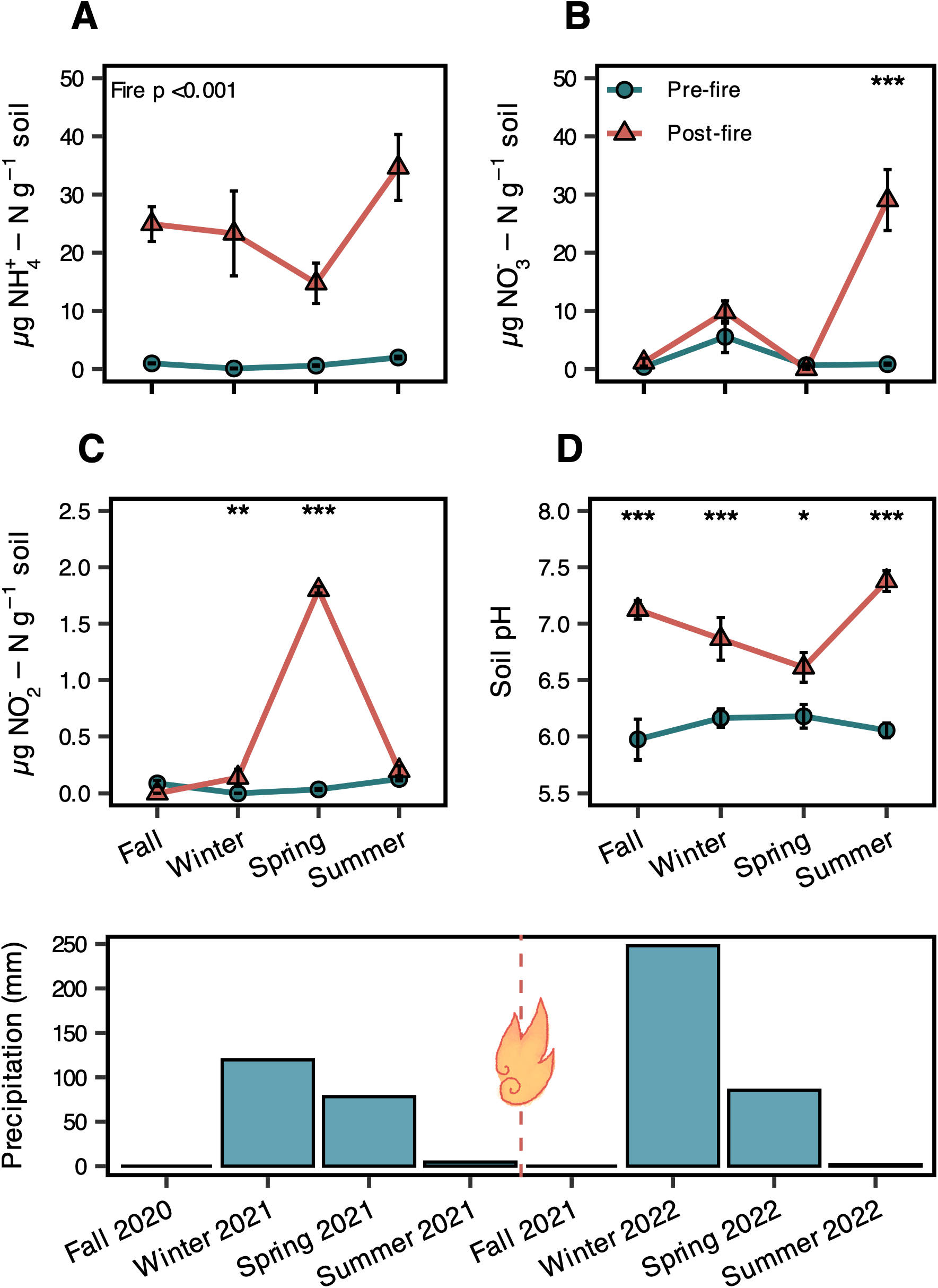
Comparison of pre- and post-fire soil NH_4_^+^ (**A**), NO_3_^-^ (**B**), NO_2_^-^ (**C**) pools and pH (**D**) across seasons. Total monthly precipitation summed over the three months preceding each sampling point is shown on the bottom panel for the entire study period, where the fire occurred in end of summer dry season 2021 demarcated by the flame symbol. “Pre-fire” (green circles) shows measurements taken over the year before the fire (Fall 2020 – Summer 2021) and “Post-fire” (orange triangles) shows measurements from same plots which we continued to sample seasonally after fire (Fall 2021 – Summer 2022). Linear mixed effects models were used to identify fire effects over time and post-hoc tests for significant differences between pre- and post-fire for individual seasons are indicated by asterisks (significance codes: ‘***’ <0.001, ‘**’ <0.01, ‘*’ <0.05). Points represent the mean and error bars are standard error (n = 5).

### 3.2 AOA and AOB amoA gene abundances

Gene copy numbers of the *amoA* gene for AOB and AOA showed a strong overall increase for bacteria, but little change in archaea in response to fire (**Figure 3**). In the year prior to the fire AOA dominated (**Figure S2**) with AOA:AOB ratios averaging 5.8 pre-fire (**Table S1**). However, AOB *amoA* gene copy numbers increased 20-fold after fire (or an average increase of 3.36× 10^6^ copy numbers g^-1^ soil over the whole year; fire × season p < 0.001; **Table S1**). Interestingly, AOB abundances did not differ from pre-fire in the first month after fire (“Fall”; **Figure 3A**), resulting in an overall post-fire AOA:AOB ratio of 1.6 (**Table S1**). However, by the winter 2022 sampling timepoint, AOB *amoA* gene abundances had increased by 22 times relative to winter the year prior to the fire (**Figure 3A**; p < 0.001), peaked during spring with an average of 54 times higher than pre-fire spring (p < 0.001), and remained elevated in summer after fire by an average of 17 times higher than pre-fire summer (**Figure 3A**; p < 0.001). AOA:AOB ratios fell to 0.65 between winter and summer after fire. In contrast, AOA *amoA* gene copy numbers did not change after fire in any season (**Figure 3B**; fire × season p = 0.2); however, there was a slight overall post-fire decrease in AOA abundance by an average of 21% or 2.67× 10^5^ copy numbers g^-1^ soil over the entire year after fire significant only at p = 0.055 (**Figure 3B**).

**Figure 3.**
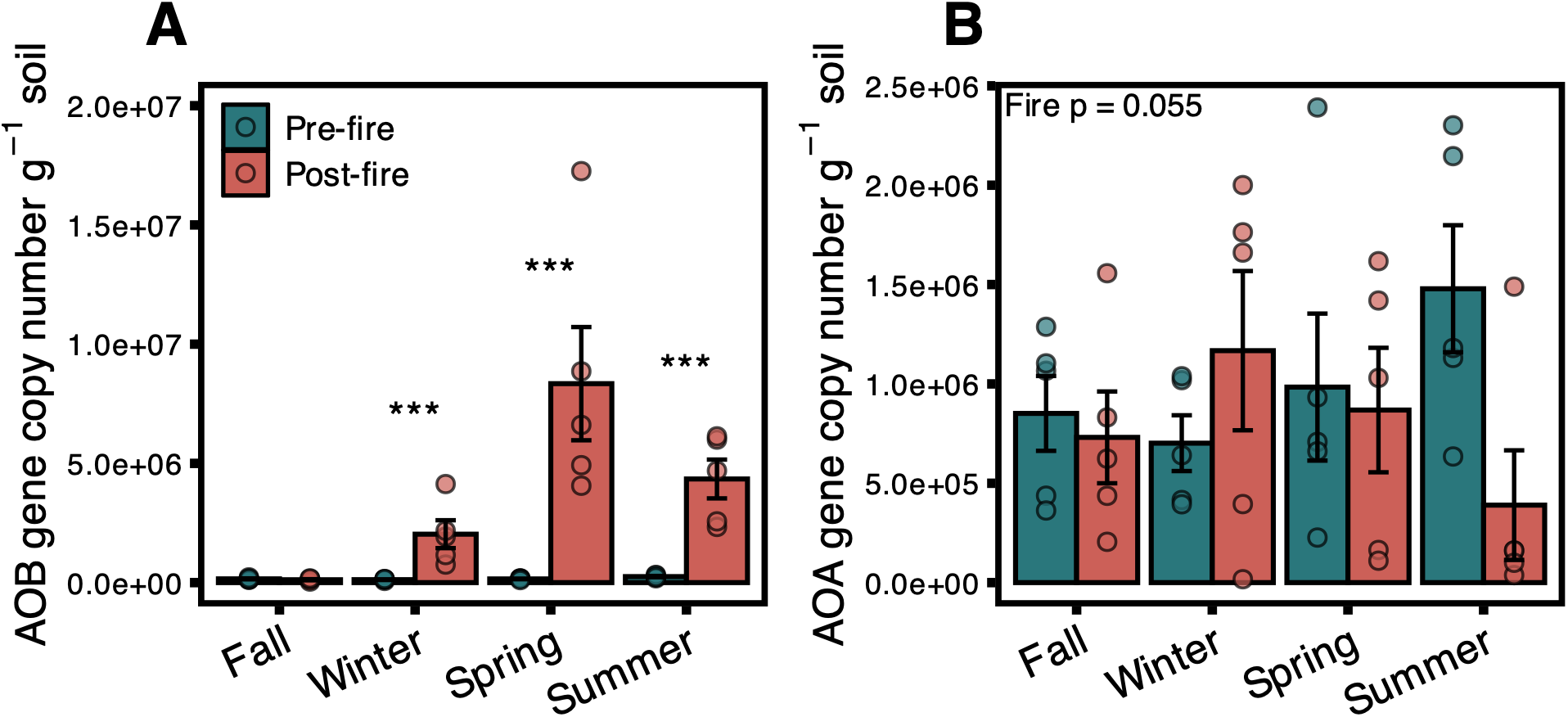
Copy numbers of the *amoA* gene in ammonia-oxidizing bacteria (AOB; **A**) and ammonia-oxidizing archaea (AOA; **B**). “Pre-fire” (green) shows the record of AOA and AOB gene abundances over the entire year before the fire (Fall 2020 – Summer 2021) and “Post-fire” (orange) shows measurements from same plots which we continued to sample seasonally after fire (Fall 2021 – Summer 2022). Linear mixed effects models were used on log-transformed variables to estimate the effect of fire across seasons and post-hoc tests for significant differences between pre- and post-fire for individual seasons are indicated by asterisks (significance codes: ‘***’ <0.001, ‘**’ <0.01, ‘*’ <0.05). Strong post-fire differences were detected for AOB in winter, spring, and summer relative to the year before fire (fire × season p <0.001; **A**), but we saw only a slight decrease in AOA after fire without a seasonal pattern (fire p = 0.055; **B**, top left). Bars represent means and error bars are standard error (n = 5).

### 3.3 Selective Inhibitions of Nitrifier Communities

Soils incubated with selective inhibition of AOA and AOB activity showed overall decreases in net N mineralization and net nitrification rates (**Figure 4**) and overall increases in N gas emissions (**Figure 5**) across treatments following fire. Post-fire net N mineralization rates decreased significantly by 99% across all treatments compared to pre-fire (**Figure 4A**; time since fire p < 0.001). Net nitrification rates decreased by 102% over all treatments following fire (**Figure 4B**; time since fire p < 0.001) driven by significant decreases in spring 2022 (post-hoc of main fire effect: p <0.001) and summer 2022 (p <0.001). There were no differences between treatments at each timepoint compared to pre-fire soils (summer 2021) for either net N mineralization rates (**A**; treatment × fire p = 0.6) or net nitrification rates (**B**; treatment × fire p = 0.3).

**Figure 4.**
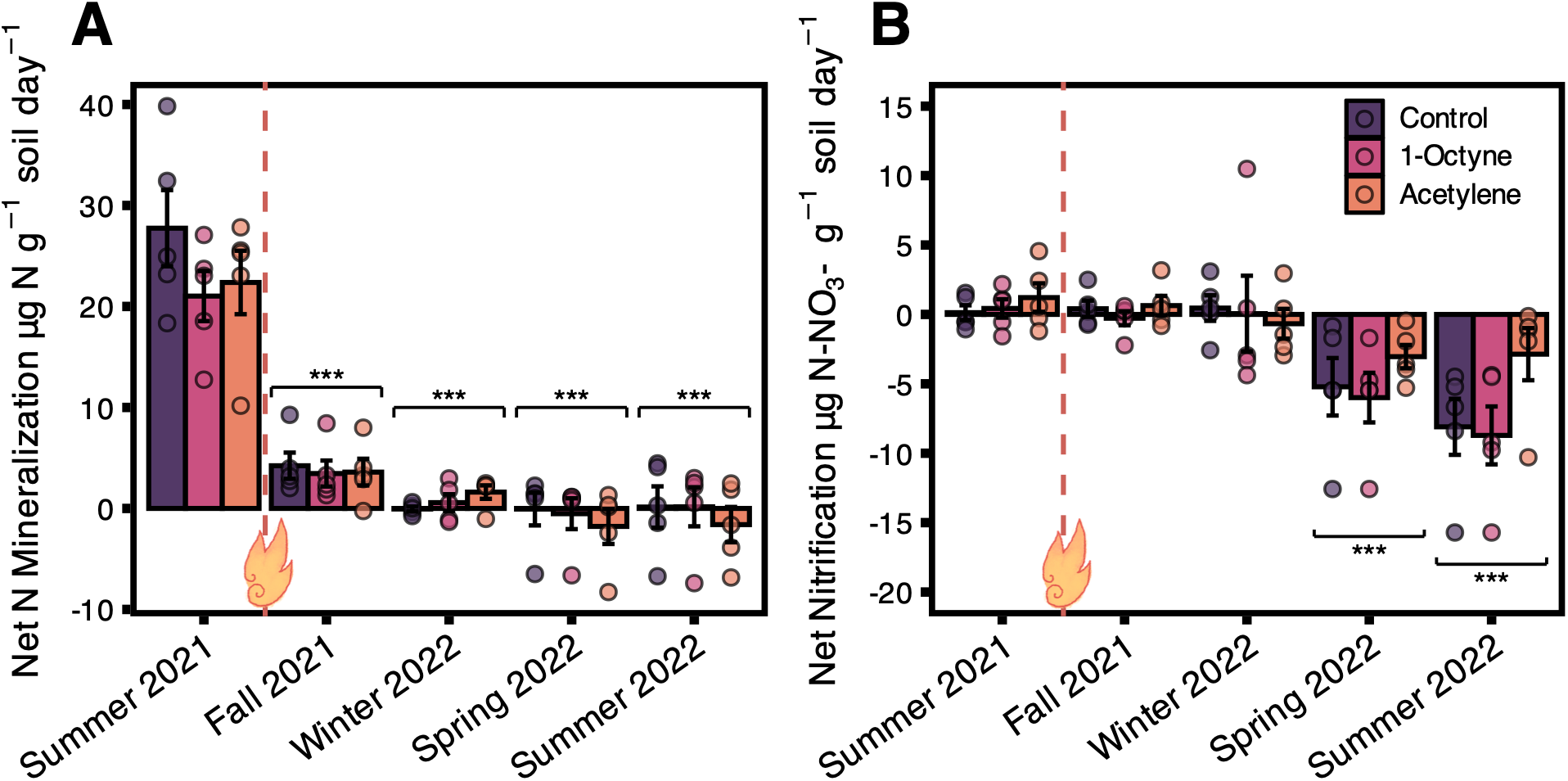
Soil net N transformation rates measured over a 72-h incubation period in the same microcosms used to measure NO and N_2_O fluxes under selective inhibitor treatments (Control, 1-Octyne, and acetylene). Summer 2021 data was collected before the site burned in an unplanned wildfire and is therefore the only pre-fire data point available. Post-fire data were collected Fall 2021 through Summer 2022 (indicated by flame symbol) and all treatments were compared to the same treatment group measured before the fire (Summer 2021) using a linear mixed effects model. Net N mineralization rates significantly decreased across all treatments post-fire (**A**; fire p < 0.001), regardless of treatment (treatment × fire p = 0.6). Similarly, time since fire mattered more than treatment for net nitrification rates (**B**; treatment × fire p = 0.3), with rates decreasing significantly post-fire in spring and fall (fire p < 0.001). P-values (‘***’ <0.001, ‘**’ <0.01, ‘*’ <0.05) indicate post-hoc investigation of the main fire effect with all treatments averaged within each sampling time compared the pre-fire measurement in Summer 2021. Error bars are standard error for each timepoint (n = 5).

**Figure 5.**
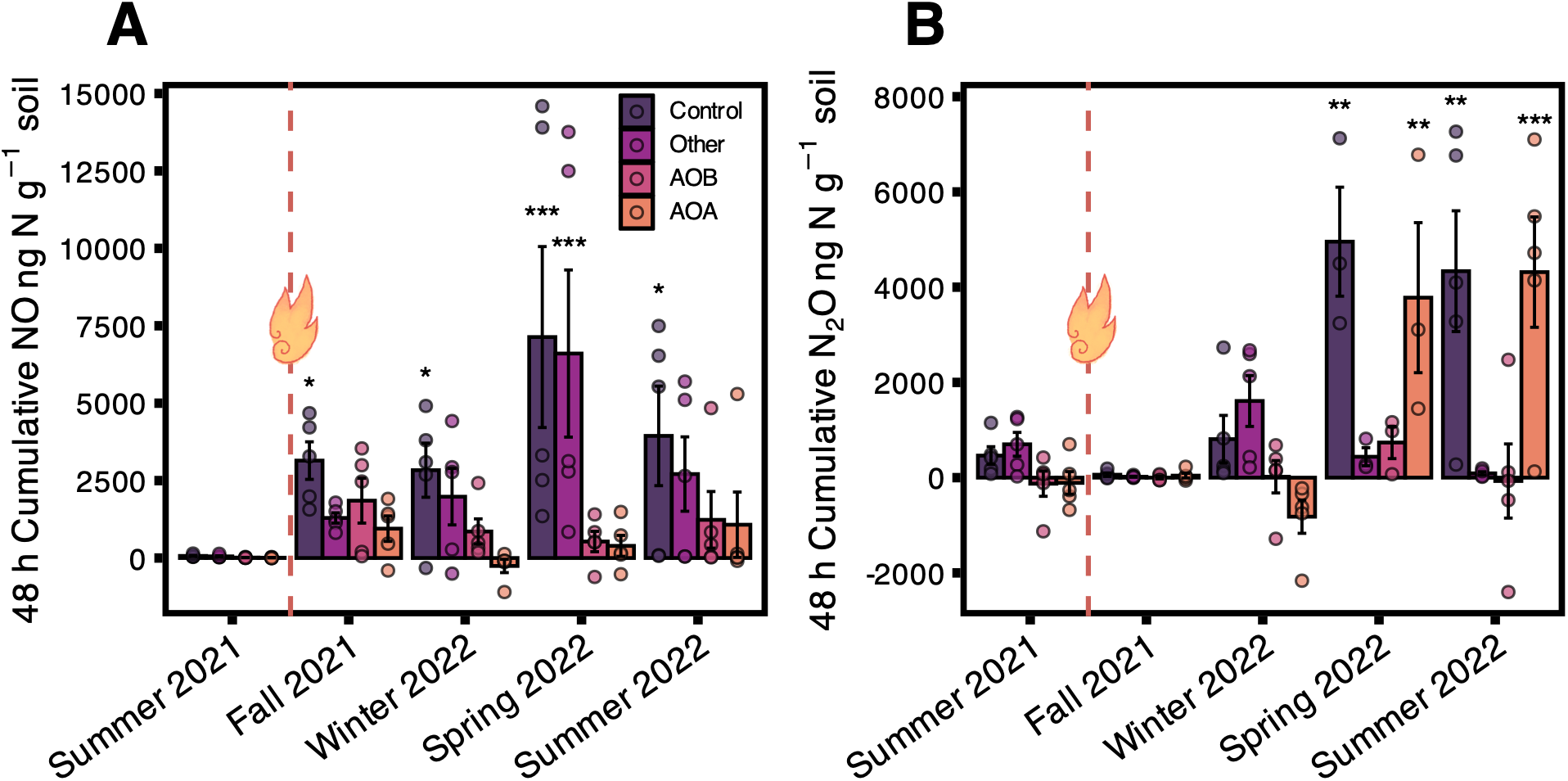
Cumulative fluxes of NO (**A**) and N_2_O (**B**) measured from soil microcosms and calculated over 48 h. Selective inhibitors were used to distinguish the contributions of AOA, AOB, and “Other” category (includes heterotrophic nitrification, abiotic, and denitrification) to soil NO and N_2_O fluxes compared to a control soil (no treatment) as described in methods section 2.3. Summer 2021 data was collected before the site burned in an unplanned wildfire and is therefore the only pre-fire data point available. Post-fire data were collected Fall 2021 through Summer 2022 (indicated by flame symbol) and all treatments were compared to the same treatment group measured before the fire (Summer 2021) using a linear mixed effects model. Significant interactions found for both NO (treatment × fire p = 0.04) and N_2_O fluxes (treatment × fire p < 0.001). P-values (‘***’ <0.001, ‘**’ <0.01, ‘*’ <0.05) show a pairwise comparison of each calculated contribution of AOA, AOB, Other, and Control groups at each post-fire sampling time to the same treatment group at the summer 2021 pre-fire timepoint. Error bars are standard error, for each timepoint and treatment n = 5, except for N_2_O in spring 2022 (**B**) where n = 3 because two replicates were lost due to an instrument communication error.

Individual contributions of AOA, AOB, Other, and Control groups to NO emissions that were calculated from selective inhibition treatments (**Figure S3**) differed over time after fire and between groups (group × time p = 0.04; **Figure 5A**). NO emissions measured from untreated soils (“Control”; **Figure 5A**) were on average 63 times higher over all seasons one year post-fire compared to pre-fire measurements in summer 2021 (p = 0.001; pre-fire mean cumulative emissions: 67 ± 19 SE ng NO-N g^-1^ soil; post-fire: 4265 ± 893 ng NO-N g^-1^ soil). AOB-derived cumulative NO emissions averaged over all seasons post-fire year were 80 times higher compared to pre-fire fluxes (p = 0.047; 13 ± 5 ng NO-N g^-1^ pre-fire, 1118 ± 314 ng NO-N g^-1^ post-fire) but did not differ significantly at any individual season due to an increase in variability after fire (p > 0.4). AOA-derived cumulative NO emissions did not differ after fire either averaged across the year (p = 0.5; 10 ± 6 ng NO-N g^-1^ soil pre-fire, 539 ± 300 ng NO-N g^-1^ soil post-fire) or in individual seasons (p > 0.9). NO emissions attributed to the “Other” category in **Figure 5A** (acetylene treatment, includes heterotrophic nitrification, denitrification, and abiotic processes) accounted for about of ∼75 % total NO emissions and were on average 59 times higher after fire (p < 0.001; 53 ± 20 ng NO-N g^-1^ soil pre-fire, 3147 ± 852 ng NO-N g^-1^ soil post-fire) driven by a significant post-fire increase in Spring 2022 (post-hoc p < 0.001). AOB-derived NO emissions did not differ from AOA-derived NO emissions within the same sampling month for any of our measurements (post-hoc p > 0.1).

N_2_O emissions derived from AOA, AOB, Other, and Control groups significantly differed over time after fire (**Figure 5B**; group × time p < 0.001). Cumulative N_2_O emissions from untreated “Control” soils averaged 462 ± 187 ng N_2_O-N g^-1^ soil at the pre-fire summer measurement and 2271 ± 634 ng N_2_O-N g^-1^ soil over one year post-fire, but varied greatly and only significantly differed from pre-fire in spring 2022 (p = 0.005) and summer 2022 (p = 0.005). AOB-derived emissions of N_2_O did not differ significantly after fire at any sampling time (p > 0.6), nor did contributions from the Other category (p > 0.9). Before the fire, AOA-derived N_2_O average cumulative emissions were very low (with inhibition treatment subtractions resulting in a negative average: −109 ± 236 ng N_2_O-N g^-1^ soil), but significantly increased to an average of 14 times higher over the entire year after the fire (1612 ± 668 ng N_2_O-N g^-1^ soil) driven by significant increases in post-fire AOA-derived N_2_O emissions in spring 2022 (p = 0.003; 3779 ± 1575 ng N_2_O-N g^-1^ soil) and summer 2022 (p <0.001; 4315 ± 1930 ng N_2_O-N g^-1^ soil).

### 3.4 Natural Abundance Isotopes of N substrates, N_2_O, and NO

Soil δ^15^N-NH_4_^+^ increased after fire only in the winter sampling (**Figure S4A**; p = 0.007) with no overall fire effect (fire p = 0.9). Combined δ^15^N-NO_3_^-^ and δ^15^N-NO_2_^-^ pools were enriched after fire in both spring and summer (**Figure S4B**; spring p = 0.004, summer p = 0.006) and significantly increased after fire overall (fire p < 0.001). Fire significantly increased δ^15^N^bulk^_N2O_ (by an average of 26‰; p < 0.001; **Figure 6**;**Figure S5A**), and site preference δ^15^N^SP^_N2O_ (by an average of 18 ‰; p = 0.03; **Figure 6**; **Figure S5B**), but did not significantly affect δ^18^O_N2O_ (p = 0.2; **Figure 6; Figure S5C**). No interactions between season and fire were detected for N_2_O isotopocules (fire × season: δ^15^N^bulk^_N2O_ p = 0.46; δ^15^N^SP^_N2O_ p = 0.3; δ^18^O_N2O_ p = 0.24; **Figure S5**). When the fractional contributions of bacterial denitrification (bD), nitrifier denitrification (nD), fungal denitrification (fD), nitrifier nitrification (Ni), and the fraction of unreduced N_2_O (r) to soil N_2_O emissions were estimated using FRAME (**Figure 7**), we found that bD and nD dominated emissions (although bD and nD were highly correlated and therefore indistinguishable from each other) but found few generalizable trends across seasons before and after fire. However, the residual N_2_O (r) fraction was lower in every season after the fire (p = 0.01; **Figure 7E**) and estimated N_2_ fluxes peaked after fire in spring and summer (**Figure 7F**). The isotopic composition of NO did not have any significant interaction with fire for δ^15^N-NO (**Figure S6A**; fire p = 0.07, fire × season p = 0.1) or δ^18^O-NO (**Figure S6B**; fire p = 0.2, fire × season p = 0.7).

**Figure 6.**
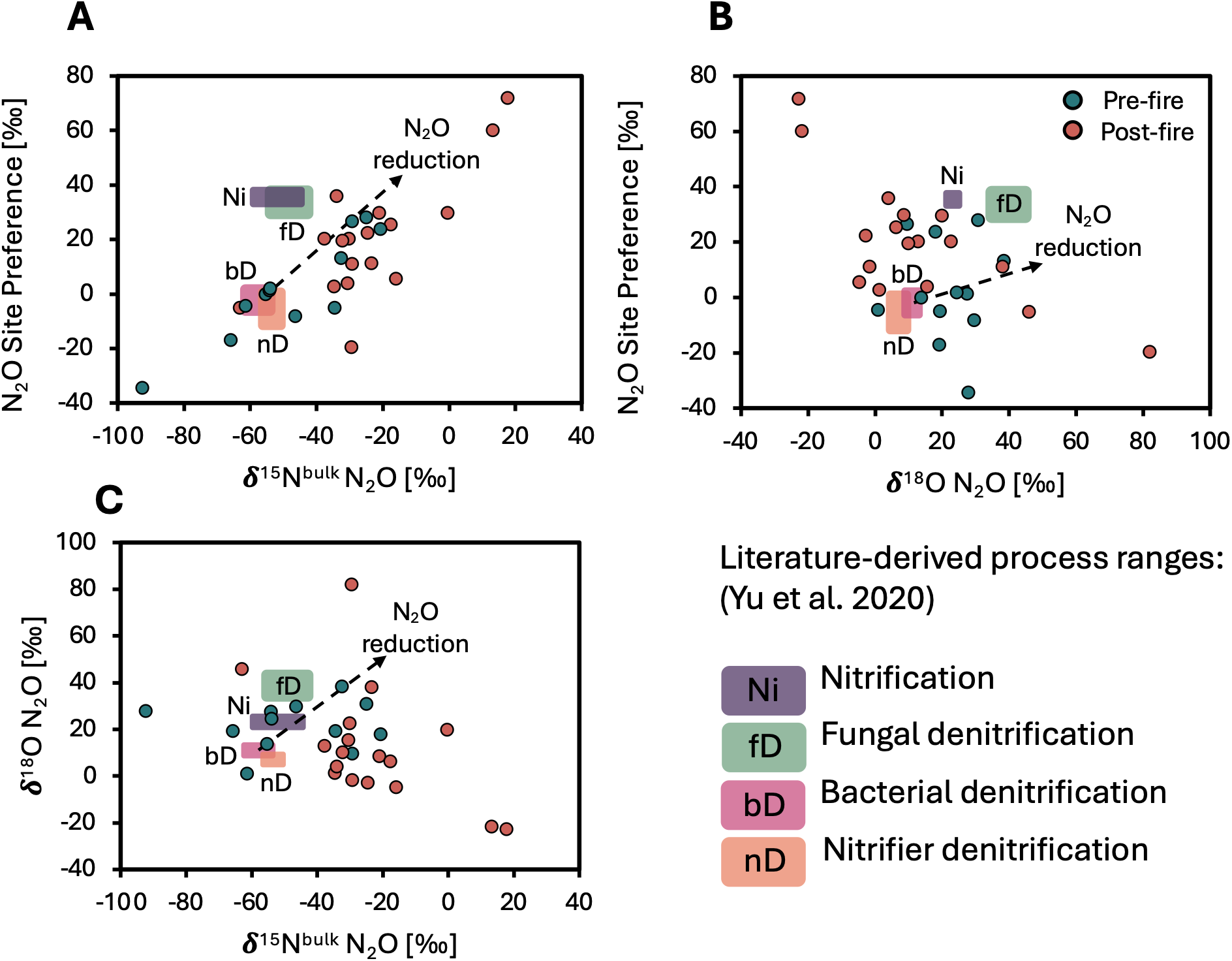
Natural abundance isotopic composition of N_2_O (δ^15^N_2_O_bulk_, δN_2_^18^O_bulk_, and site preference δ^15^N_2_O_SP_) measured from pre- and post-fire soils in 48 h incubations. Boxes indicate literature-derived ranges for nitrification (Ni), fungal denitrification (fD), bacterial denitrification (bD), and nitrifier denitrification (nD), and which were adjusted for substrate isotopic composition according to **Table S4**. “Pre-fire” (green) soils were collected over the year preceding the fire in summer 2021, spring 2021, winter 2021, and fall 2020 (n = 3 per season). “Post-fire” (orange) soils were collected over the same seasonal time points after the fire (fall 2021, winter 2022, spring 2022, summer 2022; n = 5 for each season except where a replicate was excluded due to low N_2_O production). Overall, fire significantly increased site preference δ^15^N_2_O_SP_ (by an average of 18 ‰; p = 0.03) and δ^15^N_2_O_bulk_ (by an average of 26‰; p = 0.001) but did not significantly affect δN_2_^18^O_bulk_ (p = 0.2) as shown in **Figure S5**. No seasonal effect was detected for isotopic composition of N_2_O. The black dashed arrows represent expected isotope values associated with the reduction of N_2_O to N_2_. Values that fell outside of expected ranges were included because they were not associated with any know sampling or instrument error.

**Figure 7:**
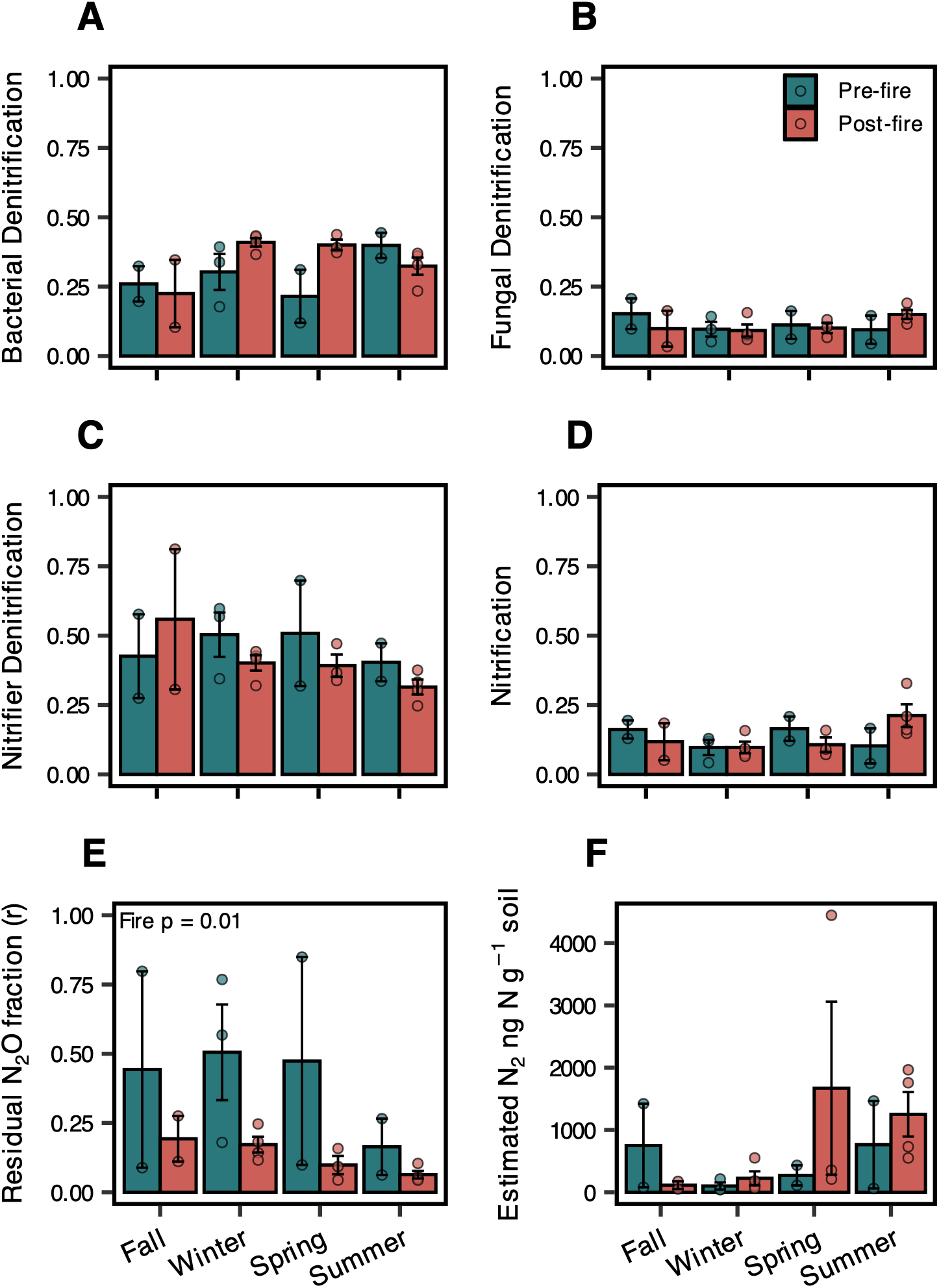
Modeled mean fractional contributions of bacterial denitrification (bD), fungal denitrification (fD), nitrifier denitrification (nD), and nitrifier nitrification (Ni) to soil N_2_O emissions (**A**-**D**) soils using the FRAME isotopic fractionation and mixing evaluation tool (https://malewick.github.io/frame/). Estimates of the fraction of unreduced N_2_O (r) are show in panel **E**, and N_2_ fluxes in panel **F** were calculated by dividing N_2_O concentrations from the residual N_2_O (r). Estimated contributions by individual soils are shown by open dots. Bars represent the mean of the individually modeled fractions with standard errors (error bars). For errors associated with each data point, see **Table S5**. There was strong correlation (> 0.7) between nD and bD in all samples, therefore it was not possible to distinguish nD from bD. Bacterial denitrification was the only microbial group that approached significant differences after fire at p = 0.057 (**A**). N_2_O (r) was consistently lower in post-fire soils (**E**; p = 0.01), indicating a greater portion of the post-fire N_2_O was reduced to N_2_, and despite lack of statistical difference, this is mirrored in the increase in peak values of estimated N_2_ flux in spring and summer (**F**).

## 4. Discussion

We used pre- and post-fire measurements to understand how wildfire promotes trade-offs in nitrifying groups and denitrifying processes that emit NO and N_2_O. As expected (**H1a**), wildfires significantly increased the abundance of AOB while leaving AOA abundance largely unchanged (**Figure 3**). However, contrary to our expectation that AOB would drive N gas emissions (**H1b; Figure 1**), we found that AOB-derived emissions did not differ significantly from pre-fire measurements; instead, overall NO emissions increased immediately after fire without dominant contributions from a single nitrifier group (**Figure 5A**). Contrary to expectations, we found a delayed post-fire increase in N_2_O emissions that was more associated with AOA than AOB (**Figure 5B**). Since AOA are not typically associated with high N_2_O production (Prosser et al., 2019), it is likely they stimulated denitrification by increasing the availability of NO_2_^-^ and NO_3_^-^ to denitrifiers. This was supported by our finding that while NO emissions increased at the first post-fire timepoints we measured, N_2_O emissions did not increase until 8 and 12 months after fire, matching timepoints where we measured peak soil extractable NO_2_^-^ and NO_3_^-^ (**Figure 2**, **Figure 4**). Additionally, changes in δ^15^N^SP^_N2O_ and δ^15^N^bulk^_N2O_ values reflected higher rates of complete N_2_O reduction to N_2_ (**Figure 6**, **Figure 7**) and provided further evidence that denitrification was stimulated at 8 and 12 months post-fire (**H2**). Together, these findings support increased nitrification activity (in particular AOA) early post-fire that supplied substrates to denitrifiers to increase NO and N_2_O emissions over time after fire.

### 4.1 Wildfire altered the abundances of AOA and AOB

Wildfire increased the abundance of AOB as hypothesized (**H1; Figure 3A**) and left AOA abundances relatively unaltered with only a slight decreasing trend that was only significant at p = 0.055 (**Figure 3B)**, consistent with other post-fire measurements (Ball et al., 2010; Jia et al., 2025; Long et al., 2014; Shu et al., 2025; Srikanthasamy et al., 2021). AOB can respond to increased NH_4_^+^ availability (Ball et al., 2010; Carey et al., 2016) and often thrive in less acidic environments where the chemical equilibrium between NH_4_^+^ and NH_3_ in soils shifts towards NH_3 (pKa_ = 9.25; Long et al., 2014; Prosser et al., 2019; Sun et al., 2024). Thus, the 30-fold increase in soil NH_4_^+^ (**Figure 2A**) and the significant increase in soil pH from 6.1 to 7.0 after fire (**Figure 2D**) could have supported AOB growth over AOA. AOA are known to respond less dramatically to increased inorganic N availabilities since AOA are better competitors in low N environments with higher NH_3_ affinities relative to AOB (Carey et al., 2016) and have also been found to decrease in abundance at high NH_4_^+^ and pH (Jia et al., 2025; Long et al., 2014; Srikanthasamy et al., 2021). Moreover, biochar, which is chemically similar to some types of wood char left behind after wildfires, has been found to increase AOB abundances with little influence on AOA (Ball et al., 2010; Sun et al., 2024), further supporting the favorability of the post-fire environment for AOB abundance.

### 4.2 NO and N_2_O emissions trade off after fire as AOA and AOB-nitrification supply substrates to denitrifiers

AOA- and AOB-derived NO emissions did not correspond directly to their abundances as we expected (**H1b**). Instead, wildfire increased post-fire variability in NO emissions across all treatments in our inhibition incubations (**Figure 5A**), obscuring clear differences between AOA and AOB contributions. It is possible that high water contents in our incubations may have limited oxygen availability for nitrification (NO is typically more associated with nitrification which is an aerobic process; Firestone & Davidson, 1989). However, NO emissions increased 60-fold over the year after fire in untreated soils (control), with significant increases in the first month after fire. As soil moisture in our incubations was identical across sampling times, this points to an active microbial community in the months immediately following fire that favored NO emissions over N_2_O. We attribute the initial increase in NO to rapid ammonia oxidation of the post-fire flush of NH_4_^+^. This mechanism is supported by the sequential accumulation of extractable soil NO_2_^-^ in spring 2022 (**Figure 2C)** and NO_3_^-^ in summer 2022 (**Figure 2B**), as well as positive net nitrification rates in fall and winter 2022 (Untreated soils only; “Control” in **Figure 4B**). Furthermore, significant ^15^N enrichment of the remaining post-fire NH_4_^+^ pool in winter 2022 **(Figure S4A**), provides strong isotopic evidence of substrate depletion during active microbial NH_4_^+^ oxidation (Deb et al., 2024).

The post-fire supply of NO_3_^-^ from nitrification likely stimulated denitrification, which may also have produced NO, particularly in Spring 2022 (**Figure 5A**) where high nitrification and denitrification activity likely co-occurred to produce the highest NO fluxes that we measured. We also found a significant portion of NO contributed by the “other” category (includes processes such as heterotrophic nitrification, abiotic processes, and denitrification; **Figure 5A**) in spring 2022, further suggesting elevated denitrification or even abiotic contributions to NO emissions, potentially promoted by the increase in NO_2_^-^. However, there were no significant differences between pre- or post-fire δ^15^N-NO or **\***δ^18^O-NO (**Figure S6**), suggesting that the underlying proportions of nitrification and denitrification activity contributing to NO emissions did not change after fire—instead, the rates of NO production from both nitrification and denitrification increased. Thus, rather than AOB driving NO emissions directly, elevated NO fluxes resulted from a broader increase in activity across all nitrifier groups including the “other” category to supply the NO_2_^-^ and NO_3_^-^ that accumulated 8-12 months post-fire.

While NO emissions immediately increased post-fire, N_2_O emissions did not differ until spring and summer 2022 (**Figure 5B**). We found no direct link between the increase in AOB abundance and N_2_O emissions; however, we found that most of the increase in N_2_O emissions we measured in spring and summer 2022 were associated with AOA activity (**Figure 5B**). This result was unexpected as AOA are associated with much lower N_2_O yields than AOB (Prosser et al., 2019). However, AOA can produce NO_3_^-^, NO_2_^-^, and hydroxylamine, which can react to form N_2_O through a variety of pathways, including abiotic processes (Giguere et al., 2017; Kozlowski et al., 2016). Indeed, we observed a significant increase in field soil-extractable NO_2_^-^ concentrations in spring 2022 (**Figure 2C**), and a ∼30-fold increase in field NO_3_^-^ concentrations in summer 2022 (**Figure 2B**), which matched timepoints with elevated N_2_O emissions associated with AOA (**Figure 5B**). Since AOA have a higher affinity for NH_3_ than AOB, they are relatively more efficient NH_3_ oxidizers (Hatzenpichler, 2012; Prosser et al., 2019) and could maintain high rates of N processing despite low abundances. Over time, the resulting accumulation of nitrifier-derived NO_2_^-^ and NO_3_^-^ could indirectly promote NO and N_2_O emissions by stimulating denitrification.

Coinciding with peak spring and summer 2022 N_2_O emissions (**Figure 5B**), we also measured significantly negative net nitrification rates (**Figure 4B**)—indicating that NO_3_^-^ was being taken up or reduced. Further, enrichment of soil δ^15^NO_3_^-^ pools (**Figure S4B**) reflects preferential enzymatic uptake of ^14^NO_3_^-^ by denitrifiers (Deb et al., 2024), and provides strong evidence for high denitrification activity. AOA can also perform well under micro-anaerobic conditions, placing them in spatial proximity to directly shuttle substrates to denitrifiers (Hatzenpichler, 2012). These converging lines of evidence suggest that despite the low AOA abundance relative to AOB, AOA communities maintained high post-fire nitrification rates, supplying intermediates such as NO_2_^-^ and NO_3_^-^ to stimulate N_2_O emissions from denitrification 8-12 months after fire. Framed within Grime’s C-S-R framework, these dynamics highlight key tradeoffs in post-fire microbial strategies. The rapid increase in AOB abundance reflects a competitor (C) / ruderal (R) strategy to acquire more post-fire resources; however, the increase in abundance did not directly translate into expected N gas flux outputs. Conversely, the relative resilience of AOA to fire suggests more thermotolerant / stress tolerant (S) traits (Goberna et al., 2012; Zhalnina et al., 2012), allowing them to maintain sustained N processing. Overall, these stress tolerant traits enabled AOA to maintain substrate supply to denitrifiers over time and shaped long-term N_2_O dynamics as post-fire resources shifted.

### 4.3 Wildfire promoted denitrification and N_2_O reduction to N_2_

We used natural abundance N_2_O isotopologues to identify major microbial groups responsible for N_2_O production, hypothesizing that their relative contributions would shift over time as microbial communities underwent post-fire successional turnover (**H2**), with an accompanying increase in nitrifier denitrification-derived N_2_O associated with increased AOB activity. Bacterial denitrification was highly correlated with nitrifier denitrification in our study and, therefore, indistinguishable in the FRAME isotope mixing models used for source apportionment (**Figure 7**). However, when considered together, both nitrifier denitrification and bacterial denitrification were the largest contributors to the total N_2_O emitted out of all processes estimated by FRAME (up to ∼50%; **Figure 6**; **Figure 7A** and **7C**). It was difficult to discern a distinct effect of fire on the contributions of fungal, bacterial, or nitrifier denitrification to N_2_O (**Figure 7**). Similar to the stability we saw in NO isotopic composition, these results suggest that postfire substrate flushes (NH_4_^+^/NO_3_^-^) accelerated rates of NO and N_2_O production without fundamentally altering the relative proportion of processes contributing to those emissions.

Despite little change observed for N_2_O-producing processes, we did find significant enrichment of δ^15^N^bulk^_N2O_ and increased δ^15^N^SP^_N2O_ in post-fire soils as would be expected of N_2_O reduction to N_2_ (**Figure S5**). While the 1 ppm headspace we used to accommodate the concentration-dependence of our instrument could have artificially increased N_2_O reduction rates, such an effect should have been uniform across all soils. Instead, we found that FRAME consistently estimated smaller residual N_2_O fractions only in wildfire-affected soils (r_N2O_; **Figure 7E**), indicating a post-fire increase in N_2_O reduction to N_2_ (as N_2_O is reduced to N_2_, the residual N_2_O substrate becomes enriched; Deb et al., 2024; Lewicka-Szczebak et al., 2020). Furthermore, when we estimated N_2_ fluxes based on isotopic composition and N_2_O concentrations in 48 h incubations, we found N_2_ emissions increased in both spring and summer after fire (**Figure 7F**), matching time points where we suspect high denitrification activity to be occurring. High rates of N_2_O reduction to N_2_ after fires have been observed in a rare few other studies, finding that N_2_ emissions were the dominant form of N gas losses in shrublands (Dannenmann et al., 2011, 2018). Increases in soil pH, like we observed (**Figure 2D**), can promote N_2_O reduction by getting closer to the pH optimum range of the N_2_O reductase enzyme (pH>7; Blum et al., 2018). There is also evidence that complete N_2_O reduction to N_2_ may be promoted by biochar (a compound chemically similar to wood char produced during wildfires), which can facilitate N_2_O reduction to N_2_ by soil denitrifiers (Cayuela et al., 2013; Tang et al., 2022; Zhang et al., 2021). The increase in N_2_O reduction to N_2_ after fire further supports the idea that high denitrification rates post-fire were promoted by intermediates generated by the activity of nitrifiers—in particular, AOA—that stimulated a diversity of denitrifying organisms aided by increased pH and wood char to increase complete reduction to N_2_.

## 5. Conclusion

Although wildfire increased AOB abundance, this did not translate into direct increases in AOB-derived NO and N_2_O emissions. Instead, post-fire N losses were governed by substrate availability (i.e., NH_4_^+^), which stimulated NO emissions across all nitrifying groups and “other” sources (such as denitrification and abiotic reactions) simultaneously. Over time, persistent AOA activity generated NO_2_^-^ and NO_3_^-^ intermediates to support N_2_O emissions post-fire via denitrification. Further, we found evidence that wildfire promoted reduction of N_2_O to N_2_, suggesting increased post-fire denitrifier activity and complete denitrification. Overall, wildfire promoted trade-offs in nitrifier community structure over time after fire, with increased nitrification activity immediately after fire driving NO emissions and handing off the substrates to support late N_2_O emissions from denitrification. As wildfire activity increases globally, resolving the temporally dynamic microbial mechanisms that govern the production of soil NO and N_2_O post-fire becomes increasingly important for understanding the implications for ecosystem N loss, air quality, and Earth’s climate.

## Acknowledgments

This research was supported by the California Department of Forestry and Fire Protection (award 8GG20812 to EZS), US Department of Agriculture (2022-67014-36675 to PMH and SIG), US Department of Energy (DE-SC0023127 to PMH and SIG), the US National Science Foundation (DEB 1916622 to PMH), and the National Science Centre Poland (Opus-2021/41/B/ST10/01045 to DLS.). Many thanks to Yareli Olazabal, Tony Calma, Melissa Zavala, Yanira (Val) Herrera, and Hoori Ajami for providing support with laboratory measurements, to Beatriz Vindiola, David Jones, and Mitchell Allen for support in the field, and to the students of BEE-BGC for their perspectives on the isotope dataset.

## CRediT author statement

**Elizah Z. Stephens**: Conceptualization, data curation, formal analysis, funding acquisition, investigation, methodology, software, validation, visualization, writing – original draft and editing. **Alexander H. Krichels**: Conceptualization, data curation, investigation, methodology, software, validation, writing – review and editing. **Aral C. Greene**: Conceptualization, data curation, investigation, methodology, validation, writing – review and editing. **Sharon Zhao**: Data curation, investigation. **Chloe Reid**: Data curation, investigation. **Maria E. Ordoñez**: Investigation, methodology, software, validation, writing – review and editing. **Jamie Irby**: Investigation, methodology. **Dominika Lewicka-Szczebak**: Conceptualization, methodology, software, validation, writing – review and editing. **Erin J. Hanan**: Conceptualization, funding acquisition, project administration, writing – review and editing. **Sydney I. Glassman**: Funding acquisition, methodology, project administration, resources, writing – review and editing. **Peter M. Homyak**: Conceptualization, funding acquisition, investigation, methodology, project administration, resources, supervision, validation, writing – review and editing.

## Data availability Statement

All data will be made publicly available in the dryad data repository.

