## Supplementary Materials for "Wildfires drive trade-offs in ammonia-oxidizing groups and promote denitrification to increase soil emissions of nitric oxide (NO) and nitrous oxide (N_2_O) in California chaparral"

Supplementary Tables:

| **Table S1** | **Pre-fire** | | | **Post-fire** | | | **Estimate** | **P-values** | | |
|---|---|---|---|---|---|---|---|---|---|---|
| **Variables (*in situ*)** | Mean | SE | n | Mean | SE | n | Fire × Season | Fire | Season | Fire × Season |
| **Clay** (%) | 17 | 0.4 | 20 | 10 | 0.4 | 20 | -7.11 | **<0.001** | 0.839 | 0.867 |
| **Sand** (%) | 63 | 0.9 | 20 | 66 | 0.4 | 20 | 2.31 | **0.013** | 0.634 | 0.686 |
| **Silt** (%) | 20 | 0.7 | 20 | 24 | 0.6 | 20 | 4.75 | **<0.001** | 0.871 | 0.899 |
| **pH** | 6.1 | 0.1 | 18 | 7.0 | 0.1 | 20 | 0.859 | **<0.001** | 0.078 | **0.009** |
| **Bulk soil C** (mg C g soil$^{-1}$) | 26.6 | 2.0 | 20 | 34.9 | 2.1 | 20 | 9.0 | **<0.001** | **<0.001** | 0.186 |
| **Bulk soil N** (mg N g soil$^{-1}$) | 1.7 | 0.1 | 20 | 2.2 | 0.1 | 20 | 0.5 | **<0.001** | **<0.001** | **0.039** |
| **C:N** | 18.7 | 0.5 | 20 | 18.4 | 0.2 | 20 | -0.3 | 0.586 | **0.027** | **0.006** |
| **Bulk soil $\delta^{15}$N** (‰) | 0.2 | 0.2 | 20 | 0.4 | 0.2 | 20 | 0.2 | 0.315 | 0.097 | 0.398 |
| **Bulk soil $\delta^{13}$C** (‰) | -27.6 | 0.1 | 20 | -26.7 | 0.3 | 20 | 0.9 | **<0.001** | **0.001** | **<0.001** |
| **Ammonium** (µg $NH_4^+$-N g soil$^{-1}$) | 0.91 | 0.17 | 20 | 27.56 | 3.55 | 20 | 26.14 | **<0.001** | **0.038** | 0.066 |
| **Nitrate** (µg $NO_3^-$-N g soil$^{-1}$) | 1.84 | 0.79 | 20 | 10.00 | 2.96 | 20 | 6.92 | **<0.001** | **<0.001** | **<0.001** |
| **Nitrite** (µg $NO_2^-$-N g soil$^{-1}$) | 0.06 | 0.01 | 20 | 0.53 | 0.17 | 20 | 0.56 | **<0.001** | **<0.001** | **<0.001** |
| **AOB** (gene copy number g$^{-1}$ soil) | $1.72\times10^5$ | $1.41\times10^4$ | 20 | $3.71\times10^6$ | $9.19\times10^5$ | 20 | $3.36\times10^6$ | **<0.001** | **0.001** | **0.001** |
| **AOA** (gene copy number g$^{-1}$ soil) | $1.00\times10^6$ | $1.41\times10^5$ | 20 | $7.89\times10^5$ | $1.57\times10^5$ | 20 | $-2.67\times10^5$ | 0.055 | 0.904 | 0.2 |
| **AOA:AOB** | 5.8 | 0.7 | 20 | 1.6 | 0.6 | 20 | -4.2 | **<0.001** | **0.056** | **0.008** |

**Table S1.** Summary of variables measured in the field over one year before fire (“Pre-fire”) and one year after fire (“Post-fire”). Means include all seasonal timepoints for both pre- and post-fire measurements. Linear mixed effect models with soil replicate as a random effect and an autocorrelation term for repeated measures over time were used to compare variables before and after fire and included an interaction for season. “Estimate” shows the average unit change in the variable after fire accounting for seasonality (positive values indicate post-fire increases; negative values post-fire decreases). P-values show significance of the effects of fire, season, and the interaction of fire and season across the entire pre- and post-fire dataset.

| Season | Pre-fire sampling date | Post-fire sampling date | Typical seasonal patterns |
|---|---|---|---|
| Fall | October 19, 2020 | October 24, 2021 | Fire season |
| Winter | February 9, 2021 | February 7, 2022 | Wet season |
| Spring | May 6, 2021 | May 19, 2022 | Plant growing season |
| Summer | August 27, 2021 | September 1, 2022 | Dry season |

**Table S2**. Seasonal sampling scheme with associated seasonal patterns typical for our chaparral site and the sampling times chosen.

| Table S3 | | | $N_2O$ (ppm) | | $NN^{15}O$ (α) | | $N^{15}NO$ (β) | | $NNO^{18}$ | |
|---|---|---|---|---|---|---|---|---|---|---|
| Replicate | Fire | Season | Mean | SD | Mean | SD | Mean | SD | Mean | SD |
| 1 | Pre-fire | Fall | 5.545 | 0.058 | 0.020257 | 2.13E-04 | 0.019686 | 1.64E-04 | 0.011384 | 7.98E-05 |
| 2 | Pre-fire | Fall | 3.279 | 0.038 | 0.011661 | 1.34E-04 | 0.011544 | 9.32E-05 | 0.006670 | 4.42E-05 |
| 3 | Pre-fire | Fall | 1.776 | 0.019 | 0.006263 | 6.77E-05 | 0.006263 | 2.99E-05 | 0.003589 | 6.87E-06 |
| 1 | Pre-fire | Winter | 1.927 | 0.017 | 0.006963 | 7.36E-05 | 0.006890 | 3.99E-05 | 0.003902 | 1.73E-05 |
| 2 | Pre-fire | Winter | 2.386 | 0.038 | 0.008693 | 1.35E-04 | 0.008523 | 9.55E-05 | 0.004875 | 4.07E-05 |
| 3 | Pre-fire | Winter | 2.088 | 0.011 | 0.007512 | 4.76E-05 | 0.007405 | 9.52E-06 | 0.004230 | 5.53E-06 |
| 1 | Pre-fire | Spring | 4.340 | 0.035 | 0.015328 | 1.22E-04 | 0.015228 | 8.60E-05 | 0.008779 | 4.42E-05 |
| 2 | Pre-fire | Spring | 2.538 | 0.038 | 0.009295 | 1.35E-04 | 0.009041 | 9.57E-05 | 0.005102 | 4.49E-05 |
| 3 | Pre-fire | Spring | 2.651 | 0.018 | 0.009344 | 6.52E-05 | 0.009347 | 2.85E-05 | 0.005362 | 1.23E-05 |
| 1 | Pre-fire | Summer | 4.287 | 0.061 | 0.015710 | 2.27E-04 | 0.015319 | 1.72E-04 | 0.008698 | 9.31E-05 |
| 2 | Pre-fire | Summer | 1.580 | 1.600 | 0.005784 | 5.79E-03 | 0.005694 | 5.66E-03 | 0.003175 | 3.19E-03 |
| 3 | Pre-fire | Summer | 1.465 | 0.017 | 0.005328 | 5.91E-05 | 0.005235 | 2.44E-05 | 0.002921 | 9.91E-06 |
| 1 | Post-fire | Fall | 2.017 | 0.016 | 0.007637 | 6.19E-05 | 0.007304 | 2.53E-05 | 0.003980 | 5.26E-06 |
| 3 | Post-fire | Fall | 1.915 | 0.016 | 0.007282 | 6.74E-05 | 0.006931 | 2.99E-05 | 0.003779 | 1.14E-05 |
| 4 | Post-fire | Fall | 1.697 | 0.014 | 0.006122 | 5.38E-05 | 0.006034 | 1.15E-05 | 0.003452 | 1.32E-06 |
| 5 | Post-fire | Fall | 1.509 | 0.023 | 0.005548 | 8.65E-05 | 0.005407 | 4.19E-05 | 0.003021 | 1.87E-05 |
| 1 | Post-fire | Winter | 1.648 | 0.019 | 0.006044 | 6.80E-05 | 0.005933 | 3.37E-05 | 0.003307 | 1.06E-05 |
| 3 | Post-fire | Winter | 1.325 | 0.009 | 0.004901 | 3.97E-05 | 0.004773 | 4.39E-06 | 0.002654 | -1.20E-06 |
| 4 | Post-fire | Winter | 2.618 | 0.056 | 0.009480 | 1.99E-04 | 0.009362 | 1.59E-04 | 0.005236 | 7.86E-05 |
| 5 | Post-fire | Winter | 3.309 | 0.082 | 0.012017 | 2.95E-04 | 0.011820 | 2.42E-04 | 0.006609 | 1.24E-04 |
| 1 | Post-fire | Spring | 8.042 | 0.146 | 0.029284 | 5.07E-04 | 0.029037 | 4.72E-04 | 0.016043 | 2.39E-04 |
| 2 | Post-fire | Spring | 2.187 | 0.030 | 0.008018 | 1.10E-04 | 0.007867 | 7.16E-05 | 0.004460 | 2.60E-05 |
| 3 | Post-fire | Spring | 1.352 | 0.011 | 0.004971 | 4.39E-05 | 0.004896 | 6.70E-06 | 0.002755 | 2.48E-06 |
| 4 | Post-fire | Spring | 2.196 | 0.030 | 0.008029 | 1.10E-04 | 0.007839 | 6.93E-05 | 0.004410 | 2.82E-05 |
| 1 | Post-fire | Summer | 4.582 | 0.058 | 0.016725 | 2.08E-04 | 0.016327 | 1.70E-04 | 0.009150 | 8.24E-05 |
| 2 | Post-fire | Summer | 2.305 | 0.036 | 0.008500 | 1.32E-04 | 0.008252 | 8.65E-05 | 0.004627 | 4.30E-05 |
| 3 | Post-fire | Summer | 3.663 | 0.048 | 0.013667 | 1.66E-04 | 0.013273 | 1.25E-04 | 0.007435 | 7.07E-05 |
| 4 | Post-fire | Summer | 3.063 | 0.035 | 0.011194 | 1.28E-04 | 0.010817 | 8.04E-05 | 0.006141 | 3.95E-05 |
| 5 | Post-fire | Summer | 4.089 | 0.059 | 0.015035 | 2.15E-04 | 0.014643 | 1.69E-04 | 0.008222 | 8.30E-05 |

**Table S3**. Isotopocule concentrations for $N_2O$ collected from soils averaged over a 3-minute measurement period (n = 180) with associated standard deviations as a measure of uncertainty. Some soils did not produce enough $N_2O$ (>1.2 ppm) to allow for isotopic analysis using our methods and so are not included here. Isotopocule concentrations were converted to delta values using equations 2-4 as displayed in Figure 6.

| Yu et al. (2020) | $\delta^{15}N_2O_{bulk}$ | | $\delta^{15}N_2O_{SP}$ | | $\delta N_2{}^{18}O$ | |
|---|---|---|---|---|---|---|
| Process | low | high | low | high | low | high |
| bD | -62.9 | -52.5 | -7.5 | 3.7 | 7.7 | 14.3 |
| nD | -57.21 | -49.61 | -13.6 | 1.9 | 3.4 | 10.3 |
| Ni | -60.51 | -43.51 | 32 | 38.7 | 20.5 | 26.5 |
| fD | -56.1 | -41.1 | 27.2 | 39.9 | 33.0 | 46.1 |
| Average ‰ of soil substrates: | $\delta^{15}N\text{-}NO_3^-$ | | $\delta^{15}N\text{-}NH_4^+$ | | $\delta^{18}O_{H2O}$ | |
| | -10.08 | | 3.49 | | -9 | |

**Table S4**. Literature-derived ranges for isotopic composition of $N_2O$ from nitrification (Ni), nitrifier denitrification (nD), bacterial denitrification (bD), and fungal denitrification (fD) from Yu et al. (2020) and Lewicki et al. (2022). Because these ranges assume a substrate contribution of 0 ‰, they were adjusted with the average combined soil $\delta^{15}N\text{-}NO_3^-$ and $\delta^{15}N\text{-}NO_2^-$ (substrates assumed to contribute to denitrification processes bD and fD) and soil $\delta^{15}N\text{-}NH_4^+$ (the substrate of Ni and nD) with the full range for these values across seasons shown in **Figure S4**. We adjusted $\delta N_2{}^{18}O$ literature ranges with the $\delta^{18}O_{H2O}$ (assumed to contribute to nD, bD, and fD) of the DI water used to wet the soils in our experiment.

| Table S5 | | | bD | | | nD | | | fD | | | Ni | | | r | | |
|---|---|---|---|---|---|---|---|---|---|---|---|---|---|---|---|---|---|
| Rep | Fire | Season | mean | CI68low | CI68up | mean | CI68low | CI68up | mean | CI68low | CI68up | mean | CI68low | CI68up | mean | CI68low | CI68up |
| 1 | Pre-fire | Fall | 0.32 | 0.10 | 0.56 | 0.27 | 0.07 | 0.48 | 0.21 | 0.05 | 0.37 | 0.19 | 0.05 | 0.36 | 0.09 | 0.04 | 0.14 |
| 2 | Pre-fire | Fall | 0.20 | 0.03 | 0.34 | 0.58 | 0.37 | 0.77 | 0.10 | 0.02 | 0.18 | 0.13 | 0.03 | 0.24 | 0.80 | 0.62 | 0.96 |
| 1 | Pre-fire | Winter | 0.34 | 0.07 | 0.63 | 0.57 | 0.29 | 0.82 | 0.05 | 0.01 | 0.10 | 0.04 | 0.01 | 0.08 | 0.57 | 0.40 | 0.73 |
| 2 | Pre-fire | Winter | 0.39 | 0.11 | 0.64 | 0.34 | 0.08 | 0.61 | 0.14 | 0.04 | 0.26 | 0.12 | 0.03 | 0.22 | 0.18 | 0.10 | 0.26 |
| 3 | Pre-fire | Winter | 0.18 | 0.03 | 0.30 | 0.60 | 0.42 | 0.77 | 0.10 | 0.02 | 0.17 | 0.13 | 0.03 | 0.23 | 0.77 | 0.56 | 0.95 |
| 1 | Pre-fire | Spring | 0.12 | 0.02 | 0.19 | 0.70 | 0.59 | 0.83 | 0.06 | 0.01 | 0.11 | 0.12 | 0.03 | 0.21 | 0.85 | 0.73 | 0.97 |
| 2 | Pre-fire | Spring | 0.31 | 0.09 | 0.54 | 0.32 | 0.10 | 0.53 | 0.16 | 0.03 | 0.29 | 0.21 | 0.06 | 0.37 | 0.10 | 0.04 | 0.15 |
| 1 | Pre-fire | Summer | 0.35 | 0.11 | 0.59 | 0.34 | 0.10 | 0.55 | 0.15 | 0.04 | 0.26 | 0.17 | 0.04 | 0.30 | 0.06 | 0.03 | 0.09 |
| 2 | Pre-fire | Summer | 0.44 | 0.11 | 0.79 | 0.47 | 0.13 | 0.79 | 0.04 | 0.01 | 0.08 | 0.04 | 0.01 | 0.07 | 0.27 | 0.16 | 0.36 |
| 4 | Post-fire | Fall | 0.10 | 0.02 | 0.16 | 0.81 | 0.75 | 0.89 | 0.03 | 0.01 | 0.06 | 0.05 | 0.01 | 0.09 | 0.11 | 0.04 | 0.17 |
| 5 | Post-fire | Fall | 0.35 | 0.11 | 0.58 | 0.31 | 0.08 | 0.53 | 0.16 | 0.04 | 0.30 | 0.18 | 0.05 | 0.33 | 0.28 | 0.16 | 0.39 |
| 1 | Post-fire | Winter | 0.43 | 0.11 | 0.72 | 0.43 | 0.13 | 0.74 | 0.07 | 0.01 | 0.13 | 0.07 | 0.01 | 0.13 | 0.18 | 0.10 | 0.26 |
| 3 | Post-fire | Winter | 0.37 | 0.13 | 0.58 | 0.32 | 0.09 | 0.56 | 0.16 | 0.03 | 0.29 | 0.16 | 0.04 | 0.29 | 0.14 | 0.07 | 0.21 |
| 4 | Post-fire | Winter | 0.43 | 0.11 | 0.76 | 0.44 | 0.14 | 0.75 | 0.06 | 0.01 | 0.11 | 0.06 | 0.01 | 0.12 | 0.25 | 0.14 | 0.36 |
| 5 | Post-fire | Winter | 0.41 | 0.10 | 0.71 | 0.42 | 0.13 | 0.70 | 0.08 | 0.02 | 0.14 | 0.09 | 0.02 | 0.17 | 0.12 | 0.06 | 0.17 |
| 1 | Post-fire | Spring | 0.39 | 0.11 | 0.69 | 0.47 | 0.18 | 0.74 | 0.07 | 0.01 | 0.12 | 0.07 | 0.02 | 0.13 | 0.04 | 0.02 | 0.07 |
| 2 | Post-fire | Spring | 0.44 | 0.13 | 0.71 | 0.37 | 0.09 | 0.63 | 0.11 | 0.02 | 0.20 | 0.09 | 0.02 | 0.16 | 0.09 | 0.05 | 0.14 |
| 3 | Post-fire | Spring | 0.37 | 0.12 | 0.63 | 0.34 | 0.09 | 0.57 | 0.13 | 0.03 | 0.25 | 0.16 | 0.04 | 0.29 | 0.16 | 0.08 | 0.23 |
| 1 | Post-fire | Summer | 0.36 | 0.11 | 0.61 | 0.38 | 0.10 | 0.62 | 0.11 | 0.02 | 0.20 | 0.15 | 0.03 | 0.27 | 0.06 | 0.02 | 0.09 |
| 2 | Post-fire | Summer | 0.33 | 0.10 | 0.55 | 0.30 | 0.08 | 0.52 | 0.16 | 0.04 | 0.30 | 0.21 | 0.05 | 0.36 | 0.05 | 0.02 | 0.08 |
| 4 | Post-fire | Summer | 0.23 | 0.06 | 0.41 | 0.25 | 0.08 | 0.42 | 0.19 | 0.04 | 0.33 | 0.33 | 0.11 | 0.54 | 0.10 | 0.04 | 0.17 |
| 5 | Post-fire | Summer | 0.37 | 0.12 | 0.62 | 0.33 | 0.09 | 0.56 | 0.14 | 0.03 | 0.25 | 0.16 | 0.04 | 0.28 | 0.04 | 0.02 | 0.07 |

**Table S5**. Modeled mean fractional contributions of bacterial denitrification (bD), nitrifier denitrification (nD), fungal denitrification (fD), nitrifier nitrification (Ni), and the fraction of unreduced $N_2O$ (r) to soil $N_2O$ emissions from selected high-emitting soils using the FRAME isotopic fractionation and mixing evaluation tool (https://malewick.github.io/frame/). Sources of $N_2O$ for each process are based on literature-derived values (Yu et al., 2020). Not all measured isotopic values resulted in reasonable model estimates and are thus not shown (compare "Rep" column, short for "Replicate", to **Table S3**). Errors associated with modeled fractional contributions for each sample are given as the upper ("CI68up") and lower ("CI68low") 68% confidence intervals.

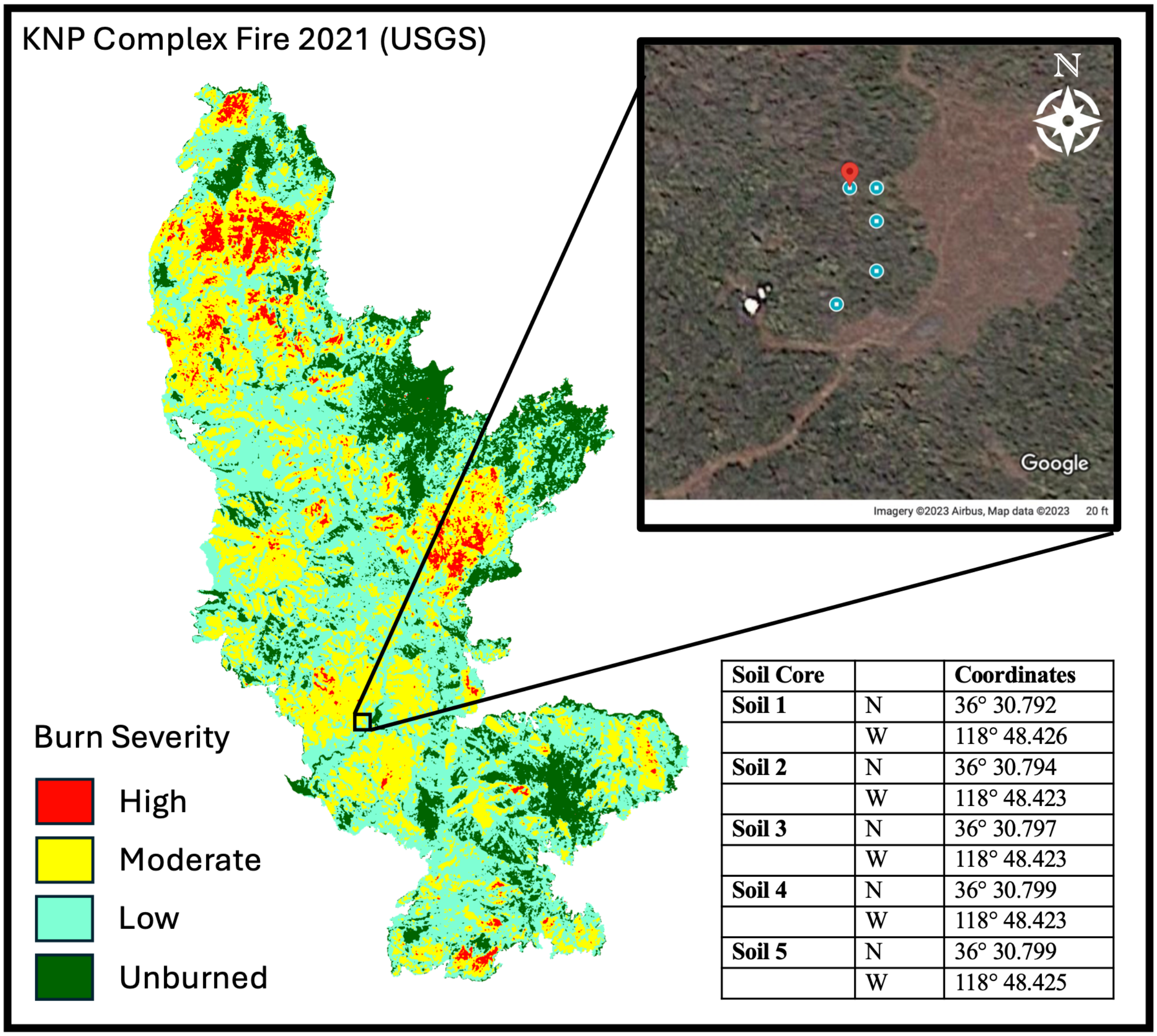


| Soil Core | | Coordinates |
|---|---|---|
| **Soil 1** | N | 36° 30.792 |
| | W | 118° 48.426 |
| **Soil 2** | N | 36° 30.794 |
| | W | 118° 48.423 |
| **Soil 3** | N | 36° 30.797 |
| | W | 118° 48.423 |
| **Soil 4** | N | 36° 30.799 |
| | W | 118° 48.423 |
| **Soil 5** | N | 36° 30.799 |
| | W | 118° 48.425 |

**Figure S1.** Burn severity of the KNP complex fire based on Burned Area Reflectance Classification from the US Geological Survey; soil burn severity at the site was classified as moderate by the Burned Area Emergency Response team (USFS BAER report). Satellite image shows transect with locations for soil core sampling which follows the contour of a small watershed. The table shows coordinates of each soil sampling location.

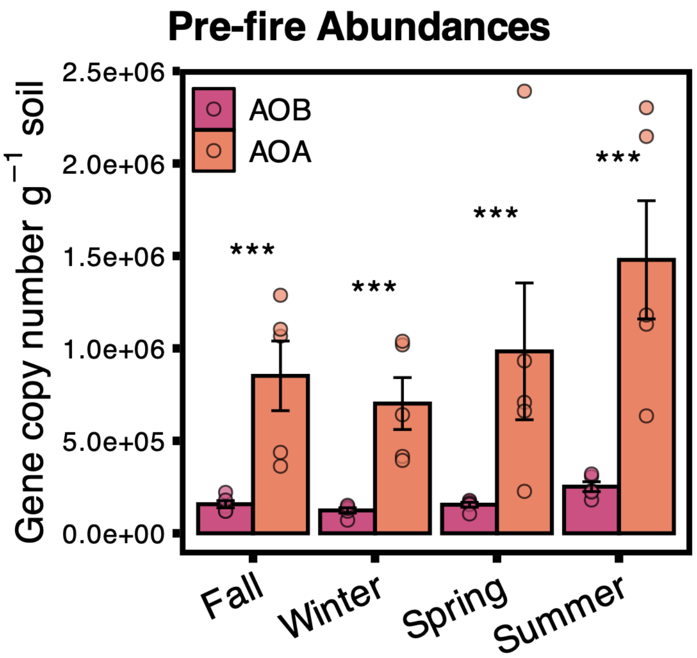


**Figure S2**. Comparison of copy numbers of the *amoA* gene in ammonia-oxidizing bacteria (AOB) and ammonia-oxidizing archaea (AOA) over one year before the fire (Fall 2020 – Summer 2021). Linear mixed effects models were used on log-transformed variables to estimate differences in AOA/AOB abundances in each season. AOA were significantly more abundant than AOB in all seasons before the fire with p-values for differences in individual seasons shown with asterisks (‘***’ <0.001, ‘**’ <0.01, ‘*’ <0.05). Bars represent means and error bars are standard error (n = 5).

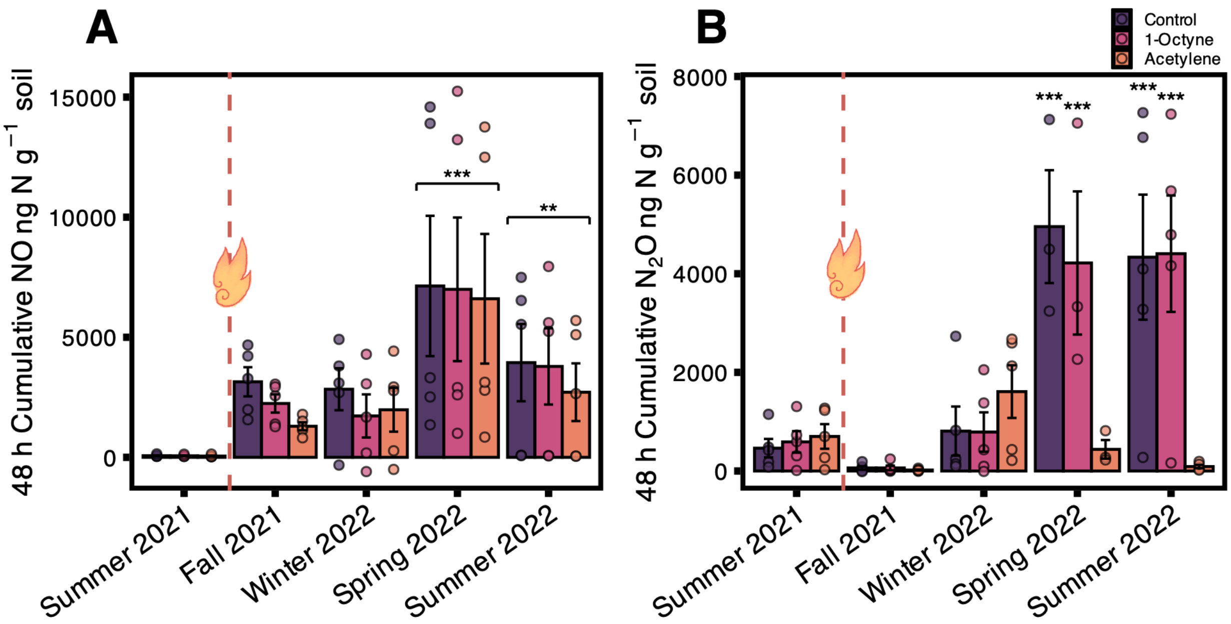


**Figure S3.** Cumulative fluxes of NO (**A**) and $N_2O$ (**B**) calculated over 48 h from soils treated with either no inhibitor ("Control", added water only), 1-octyne to selectively inhibit AOB activity ("1-Octyne"), or acetylene to inhibit both AOA and AOB ("Acetylene"). Post-fire (Oct 2021 through Sep 2022) treatments were compared to the same treatment group measured before the fire (Sep 2021) using a linear mixed effects model. Fire did not affect individual treatments significantly for NO emissions (**A**; treatment × fire $p = 0.9$), but NO emissions increased from all treatments after fire (**A**; fire $p < 0.001$). Therefore, in **A**, p-values ('***' <0.001, '**' <0.01, '*' <0.05) indicate post-hoc comparison of main fire effect with all treatments averaged over each sampling time compared to the pre-fire measurements in summer 2021. For $N_2O$, individual treatments interacted significantly with fire (**B**; treatment × fire $p < 0.001$), and p-values in **B** show a pairwise comparison of post-fire treatment groups at each sampling time to pre-fire measurements in summer 2021. Calculated contributions from each inhibited group are shown in in **Figure 4**. Error bars are standard error, for each timepoint n = 5.

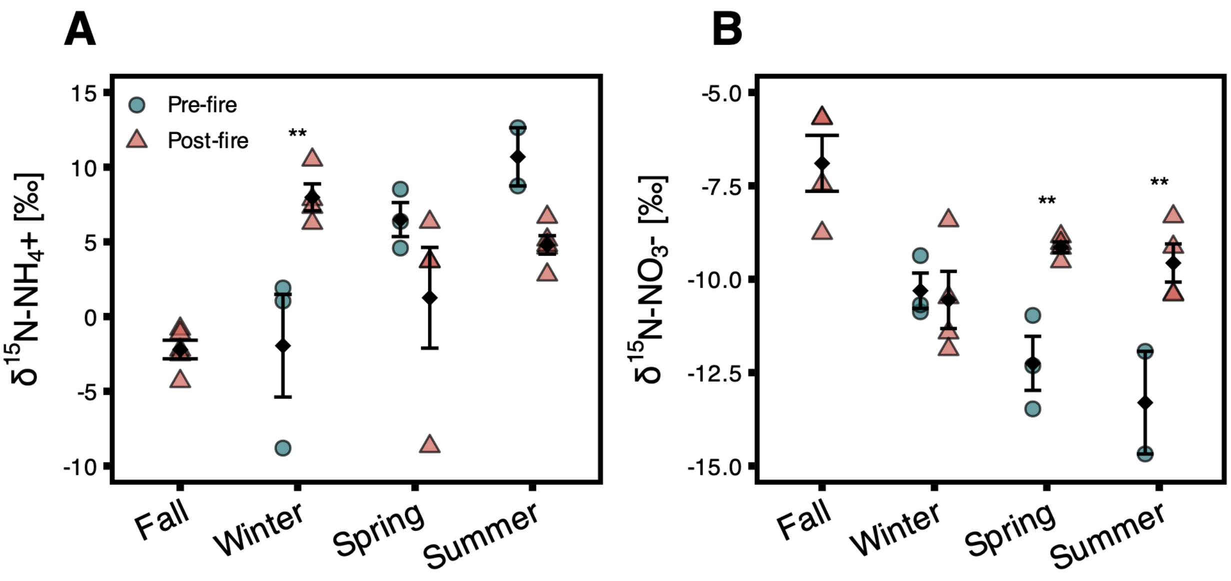


**Figure S4**. Isotopic composition of soil extractable δ$^{15}$N-$NH_4^+$ (**A**) and extractable δ$^{15}$N-$NO_3^-$ + δ$^{15}$N-$NO_2^-$ (**B**). Black diamonds represent means and error bars are standard errors, for each timepoint n = 3 pre-fire, n = 5 post-fire (samples from fall pre-fire failed due to low extraction concentrations and are therefore not included). Fire significantly interacted with season for both δ$^{15}$N-$NH_4^+$ ($p = 0.005$) and δ$^{15}$N-$NO_3^-$ ($p = 0.02$) with individual differences between pre- and post-fire measurements in each season represented by '***' <0.001, '**' <0.01, '*' <0.05 (post-hoc comparison).

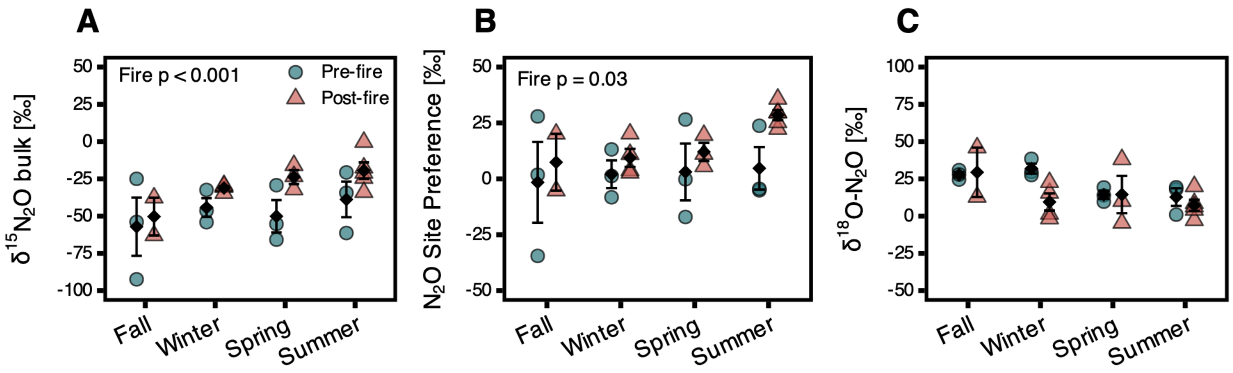


**Figure S5**. Natural abundance of $N_2O$ isotopes presented seasonally over the study period. “Pre-fire” (green) soils represent samples taken over the year preceding the fire in summer 2021, spring 2021, winter 2021, and fall 2020 (n = 3 per season). “Post-fire” (orange) soils were sampled over the same seasonal time points after the fire (fall 2021, winter 2022, spring 2022, summer 2022; n = 5 for each season except where a replicate was excluded due to low $N_2O$ production). We used a linear mixed effects model with interactions terms for fire and season to test for differences between pre- and post-fire measurements (p-values in top left). Overall, fire significantly increased $\delta^{15}N^{bulk}{}_{N2O}$ by an average of 26‰ (**A**; fire $p < 0.001$), and site preference $\delta^{15}N^{SP}{}_{N2O}$ by an average of 18‰ (**B**; fire $p = 0.03$), but did not significantly affect $\delta N_2{}^{18}O_{bulk}$ (**C**; fire $p = 0.2$). No significant seasonal effect was found for any isotopomer (fire × season: **A**, $p = 0.46$; **B**, $p = 0.3$; **C**, $p = 0.24$). Black diamonds represent means and error bars are standard error, for each timepoint n = 5 except where too little $N_2O$ production prevented isotopic characterization.

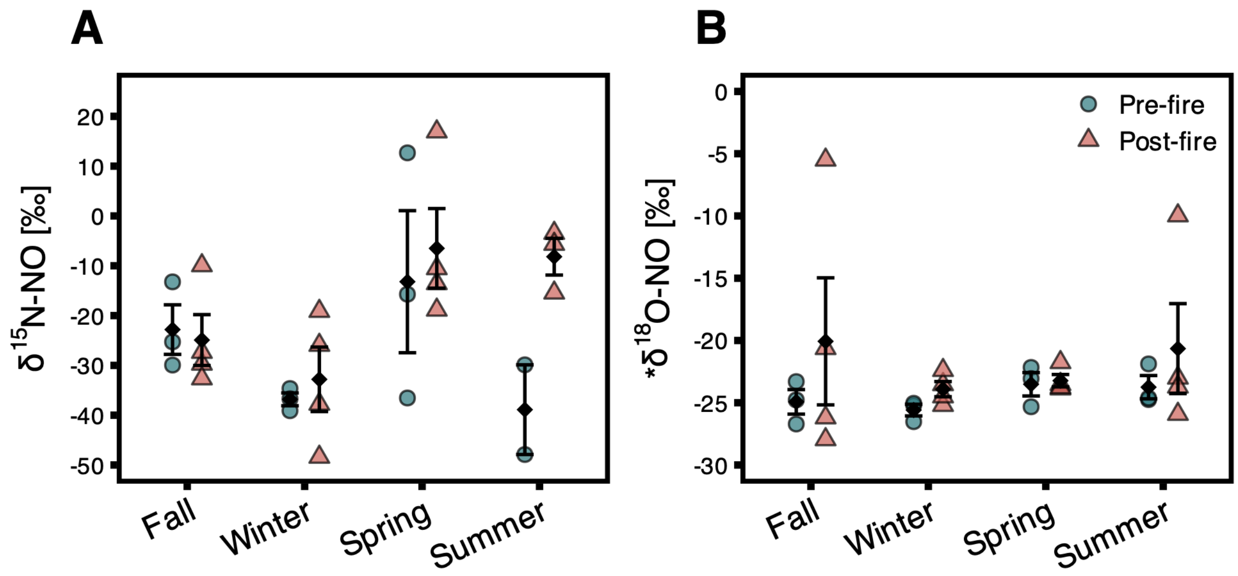


**Figure S6**. Isotopic composition of NO collected over 72-h incubations in sealed jars. The asterisk symbol indicates that *$\delta^{18}$O-NO values are uncorrected and therefore not the true $\delta^{18}$O-NO values as the oxygens in nitrite are known to exchange with oxygen in the water used to extract nitrite from the NOx pads and are therefore roughly 25‰ lower than the true values (see Methods S2). However, any exchange of oxygen between nitrite and water should have occurred uniformly across samples, allowing us to assess potential differences in NO-producing processes pre and post fire. No significant interaction with fire was found for $\delta^{15}$N-NO (fire $p = 0.07$, fire × season $p = 0.1$) or *$\delta^{18}$O-NO (fire $p = 0.2$, fire × season $p = 0.7$). Black diamonds represent means and error bars represent standard errors ($n = 5$), except where extractions yielded too little $NO_2^-$ for analysis.

**Supplementary Methods**

***Methods S1. Substrate ($NH_4^+$ / $NO_3^-$) isotopic characterization***

To reference our sample isotopic values to previously published ranges of isotopologues of $N_2O$ for distinct microbial processes (nitrification, nitrifier denitrification, bacterial denitrification, and fungal denitrification), we adjusted these values using the average combined soil $\delta^{15}N$-$NO_3^-$ and $\delta^{15}N$-$NO_2^-$ (substrates assumed to contribute to bacterial and fungal denitrification processes), $\delta^{15}N$-$NH_4^+$ (substrate assumed to contribute to nitrification and nitrifier denitrification) and $\delta^{18}O_{H2O}$ according to recommendations by Yu et al. (2020; **Table S4**). $\delta^{15}N$-$NH_4^+$ was characterized using acid trap diffusion (Brooks et al., 1989; Irby et al., 2026), where $NH_4^+$ is volatilized to $NH_3$ by increasing the pH with magnesium oxide, then the $NH_3$ is collected on a paper disk acidified with 2.5 M potassium bisulfate. $\delta^{15}N$-$NO_3^-$ was similarly quantified by addition of Devarda's alloy to reduce $NO_3^-$ to $NH_4^+$ as a secondary step after volatilizing any present $NH_4^+$. Paper disks were desiccated before analysis and measured on a Costech elemental analyzer (Costech Elemental Combustion System) coupled to an isotope ratio mass spectrometer (Delta V Advantage IRMS, $\delta^{15}N$ precision = 0.06 ‰) at FIRMS. $\delta^{18}O_{H2O}$ of deionized water in the building was measured with a Triple Isotopic Water Analyzer (TIWA-DLT-EP, Model 912-0032). Final adjustments to literature-derived process ranges for isotopes of $N_2O$ accounting for soil substrates were: -10.1 ± 2.1 ‰ for combined $\delta^{15}N$-$NO_3^-$ and $\delta^{15}N$-$NO_2^-$, 3.5 ± 5.0 ‰ for $\delta^{15}N$-$NH_4^+$, and -9.0‰ for $\delta^{18}O_{H2O}$ (**Table S4**).

***Methods S2. Isotopic composition of NO***

To characterize the isotopic composition of NO emitted from soils, jar lids were fitted with $NO_x$ precoated collection pads (Ogawa & Co., Pompano Beach, FL, USA) to capture NO as described in Homyak et al. (2016). The pads were installed at the beginning of the same incubation described above for isotopic $N_2O$ characterization. Pads were inserted immediately after wet-up and immediately sealed in the jars before the flushing step to minimize exposure to outside air. At the end of the 24 h incubation, pads were removed and NO captured in the pads was extracted as $NO_2^-$ in DI water and frozen at -20 °C before being analyzed for $\delta^{18}$O-NO and $\delta^{15}$N-NO using the bacterial denitrifier method (Sigman et al., 2001) at the Facility for Isotope Ratio Mass Spectrometry (FIRMS, https://ccb.ucr.edu/facilities/firms) at the University of California, Riverside. *Pseudomonas aureofaciens* were used to convert $NO_2$- in the extracts to $N_2O$ and $\delta^{15}$N and $\delta^{18}$O values were measured using a Thermo Delta V isotope ratio mass spectrometer (Thermo Fisher Scientific) with international reference materials USGS-32, USGS-34, and USGS-35 included in each analytical run. These values are reported as *$\delta^{18}$O-NO because they are not the true $\delta^{18}$O-NO values as 1) the oxygens in nitrite are known to exchange with oxygen in the water used to extract nitrite from the NOx pads (Casciotti et al., 2007; Dahal & Hastings, 2016), and 2) $\delta^{18}$O-NO values must be further corrected to account for the loss of less oxygen in the conversion from nitrite samples to $N_2O$ compared to nitrate to $N_2O$ (Casciotti et al., 2007), thus, *$\delta^{18}$O-NO values are roughly 25‰ lower than the true values (Dahal & Hastings, 2016). However, any exchange of oxygen between nitrite and water should have occurred uniformly across samples, preserving a $\delta^{18}$O-NO signal from the NOx collection pads (Coughlin et al., 2017) and allowing us to assess potential differences in NO-producing processes pre and post fire (e.g., a post-fire shift from hydroxylamine oxidation to more nitrifier denitrification would have altered *$\delta^{18}$O-NO because of differences in oxygens derived from air vs. water; Homyak et al., 2016).
